# Innexin-mediated adhesion between glia is required for axon ensheathment in the peripheral nervous system

**DOI:** 10.1101/2022.07.12.499769

**Authors:** Mriga Das, Duo Cheng, Till Matzat, Vanessa J. Auld

## Abstract

Glia are essential to protecting and enabling nervous system function and a key glial function is the formation of the glial sheath around peripheral axons. Each peripheral nerve in the Drosophila larva is ensheathed by three glial layers, which structurally support and insulate the peripheral axons. How peripheral glia communicate with each other and between layers is not well established and we investigated the role of Innexins in mediating glial function in the Drosophila periphery. Of the eight Drosophila Innexins, we found two (Inx1 and Inx2) are important for peripheral glia development. In particular loss of Inx1 and Inx2 resulted in defects in the wrapping glia leading to disruption of the glia wrap. Of interest loss of Inx2 in the subperineurial glia also resulted in defects in the neighbouring wrapping glia. Inx plaques were observed between the subperineurial glia and the wrapping glia suggesting that gap junctions link these two glial cell types. We found Inx2 is key to Ca+2 pulses in the peripheral subperineurial glia but not in the wrapping glia, and we found no evidence of gap junction communication between subperineurial and wrapping glia. Rather we have clear evidence that Inx2 plays an adhesive and channel-independent role between the subperineurial and wrapping glia to ensure the integrity of the glial wrap.

## INTRODUCTION

Glial cells are required for the development and maintenance of the nervous system and provide multiple functions in this role, a key function being insulation or ensheathment of peripheral nerves. In the peripheral nervous system (PNS) glia ensheath axons, provide insulation and participate in the formation of the blood-nerve barrier (Carlson et al., 2000). In vertebrates, two types of Schwann cells, myelinating and non-myelinating wrap and insulate peripheral axons. The former makes a multilayered myelin sheath around large caliber axons, whereas the latter directly ensheath small caliber axons to form Remak bundles. Myelinating Schwann cells (SC) have been the focus of vertebrate literature and dye diffusion studies in the myelin sheath found that low molecular mass compounds can diffuse between the inner and perinuclear SC cytoplasm via gap junctions (Cina et al., 2007). The role of gap junctions in non-myelinating Schwann cells (NMSC), however remains unknown.

In *Drosophila*, the role of nerve ensheathment is performed by three glial layers: perineurial glia (PG), subperineurial glia (SPG) and wrapping glia (WG). The outermost glial layer formed by the PG, contacts and is covered by the neural lamella (an extensive extracellular matrix). The intermediate glial layer, the SPG, encircle the entire nerve bundle and form autocellular septate junctions which are the structural basis for the blood-nerve-barrier (BNB). The innermost glial layer, the WG, directly ensheath axons and separate them into bundles that resemble vertebrate Remak bundles formed by vertebrate NMSCs. The mechanisms underlying adhesion or communication between these glial layers are poorly understood.

Gap junctions are formed when a hemichannel pairs with a hemichannel from an apposing cell membrane. These channels facilitate direct cell-cell communication by allowing ions and small molecules to pass through them. In addition, vertebrate connexin-based gap junctions can function as cell adhesion proteins in a channel independent manner (Bruzzone et al., 1996; Kumar and Gilula, 1996; Phelan et al., 1998; Elias et al., 2007). Gap junctions in invertebrates are composed of innexins and eight innexin genes have been identified in *Drosophila melanogaster* (Bauer et al., 2005). Although innexins and connexins do not share sequence similarity, they perform similar functions (Skerrett and Williams, 2016). For example, propagation of calcium waves in astrocytes is primarily facilitated by connexins (Scemes and Giaume, 2006). Similarly, Speder and Brand (2014) found that calcium oscillations in the SPG of the *Drosophila* CNS are gap junction mediated, as are Ca+2 oscillations in Drosophila perineurial glia (Weiss et al., 2022). However, the role of innexins in mediating glial communication between the three glial layers of the Drosophila peripheral nerve remains unknown.

We set out to identify which gap junction proteins are present and required in the peripheral glia and determined that Inx1 and Inx2 are expressed throughout the peripheral glia and in particular between the SPG and WG membranes. In the WG, loss of both Inx1 and Inx2 lead to reduced axonal ensheathment and in extreme cases fragmentation of the WG. Moreover, knockdown of Inx2 in the SPG did not affect the SPG but lead to a cell non-autonomous effect on the WG. We show that calcium pulses in the SPG are mediated by Inx2 based gap junctions between SPG, but not in the WG nor between the SPG and WG, suggesting that Inx function between these two glial layers is not based on channel function. Indeed we determined that Inx2 functions in an channel-independent manner likely as an adhesion protein to mediated the interactions between the SPG and the WG.

## MATERIALS AND METHODS

### Fly strains and genetics

The following fly strains were used in this study: repo-GAL4 (Sepp et al., 2001); Nrv2-GAL4 (Sun et al., 1999), 46F-GAL4 (Xie and Auld, 2011); Gli-GAL4 (Sepp and Auld, 1999); SPG-GAL4 (Schwabe et al., 2005); UAS-mCD8::GFP (Lee and Luo, 1999); UAS-Dicer2 (Dietzl et al., 2007); UAS-mCD8::RFP (Bloomington Stock Center, BDSC); UAS-RFP::Inx2 (Speder and Brand, 2014); UAS-R-Inx2-RFP, UAS-R-Inx2[L35W]-RFP, UAS-R-Inx2[C256S]-RFP (Miao et al., 2020); UAS-p35 (BDSC); UAS-GCaMP6S (BDSC) (Tian et al., 2009). The following GFP protein-trap insertions were used: Nrv2::GFP, Jupiter::GFP and NrxIV::GFP (Morin et al., 2001) (Kelso et al., 2004); (Buszczak et al., 2007). The following RNAi lines were used: Inx2-RNAi - JF02446 (BDSC), KK111067 (Vienna Drosophila Resource Center, VDRC), 4590R-3 (National Institute of Genetics, NIG); Inx1-RNAi - GD3264 (VDRC); Inx3-RNAi - GD14965 (VDRC), HM05245 (BDSC); Inx7-RNAi - KK112684 (VDRC); Atg1-RNAi – GD7149, VSH330433 (VDRC); Atg18-RNAi –KK100064, GD12342 (VDRC). All RNAi experiments were carried out at 25°C without Dicer2 in the background unless specified. All GAL4 controls were crossed to *w*^*1118*^.

### Immunolabeling and image analysis

Larvae were dissected and fixed for immunolabeling using previously described methods (Sepp et al., 2000). The following primary antibodies were used in this study: guinea pig anti-Inx2 (1:500), (Smendziuk et al., 2015); rabbit anti-Inx1 (1:50), (Bauer et al., 2004); rabbit anti-HRP (1:500, Jackson ImmunoResearch, West Grove, PA); mouse anti-Futsch/22C10 (1:1000, DSHB); rabbit anti-p35 (1:1000, Novus Biologicals, Oakville, Ca); rabbit anti-Dcp1 (1:1000, Cell Signaling Technology). The following secondary antibodies were used at a 1:300 dilution: goat anti-mouse Alexa 488, Alexa 568 and Alexa 647; goat anti-rabbit Alexa 568 and goat anti-guinea pig Alexa 647 (Molecular Probes). DAPI (1:1000, Invitrogen) was used to stain nuclei. Images were obtained using Delta Vision Spectris (Applied Precision/GE Healthcare) using a 60x oil immersion objective (NA 1.4). An image was captured every 0.2 μm and the resulting stacks were deconvolved (SoftWorx, Applied Precision/GE Healthcare) using a point spread function measured with 0.2 μm beads conjugated to Alexa dyes (Molecular Probes) and mounted in Vectashield (Vector Laboratories, Burlington, Canada). Orthogonal sections were generated using SoftWorx. A single z-slice, conveying the information relevant to the experiment, was chosen from each z-stack and images were compiled using Adobe Photoshop and Adobe Illustrator CC. For transmission electron microscopy analysis larvae were dissected and prepared using previously described methods (Matzat et al., 2015).

### Larval tracking

For each larval tracking session, multiple 3_rd_ instar larvae were added to a fresh 2% agar plate. Food-safe dye was added to enhance contrast. Larval movements were recorded continuously for 60 seconds using a Canon VIXIA HF R800 video camera (Canon). The recorded movies were analyzed using the Fiji (ImageJ) plug-in wrmTrck (Brooks et al., 2016) to calculate average speed and travel distance. At least two replicates were carried out for each genotype.

### *In vivo* calcium imaging

*UAS-GCaMP6S* was expressed in the SPG and WG using *Gli*-*GAL4* and *Nrv2-GAL4* respectively. Third instar larvae were anesthetized using isoflurane for 4 mins, on average. Each larva to be anesthetized was placed in a 50ml tube containing a Kimwipe soaked with 300μl of isoflurane. The larva was removed from the tube when visible movement had ceased (approximately 2-4mins). Each larva was placed ventral side up on a prepared agarose slide and gently pressed with a 18×18 mm coverslip and tape to reduce movement. GCaMP6S fluorescence was imaged using a Leica SP5 II laser scanning confocal microscope with a tandem scanner and HyD detector. Image stacks of the posterior region of the ventral nerve cord and peripheral nerves were collected using a 25X water objective (NA 0.95). 3D projections of the stacks were acquired using the Leica Application Suite Advanced Fluorescence (Leica AF software). Each image was acquired at a speed of 8,000 lines per second with a line average of 4 resulting in a collection time of 131 ms per frame at a resolution of 512 × 512 pixels. The pinhole was opened to 2.5-4.5 Airy units (AU). The total z-steps taken for each stack were set to around 25-52 steps. ROIs were manually selected and the mean fluorescence intensity was measured using the Leica AF software and were plotted.

### Statistics

Prism (GraphPad Software) was used for all statistics analysis and a One-way Anova utilized to compared multiple genotypes with the specified control with a Dunnet multiple comparison test. For all image analyses at least 6-7 nerves were analyzed per larvae.

## RESULTS

### Innexin 1 and Innexin 2 are expressed in glial cells of the larval peripheral nerve

To test for the presence of innexins in the peripheral glia, we first used an RNAi approach to knockdown those innexins known to be expressed in Drosophila glia (Inx1, 2, 3, 7)(Stebbings et al., 2002; Ostrowski et al., 2008). RNAi-mediated knockdown in all glia using the pan glial driver, *repo-GAL4*, generated peripheral glia phenotypes for only Inx1 and 2 (Table 1). Pan glial knockdown of Inx3 and Inx7 did not affect glial or nerve morphology (Table 1). We thus examined the distribution of Inx1 and Inx2 in the peripheral glia in 3^rd^ instar larvae. At this stage, each peripheral nerve is surrounded by three glial layers, the innermost wrapping glia (WG) that wrap the axons, the intermediate subperineurial glia (SPG) that form the blood-nerve barrier and the outermost perineurial glia (PG). A combination of glial layer specific markers were used in conjunction with antibodies specific to Inx1 (Bauer et al., 2004) and Inx2 (Smendziuk et al., 2015). We first assayed the distribution of Inx2 in nerves where the PG were labeled with Jupiter::GFP along with a membrane tagged RFP in the SPG (*SPG-GAL4*>*UAS-mCD8::RFP)* (Fig. 1A). Inx2 immunolabeling was observed as puncta that associated with the PG as well as the SPG membranes (Fig. 1A and B, arrowheads). Inx2 positive puncta were also observed at the center of the nerve where the axons and WG normally reside (Fig. 1A and B, arrowheads). The presence of Inx2 positive puncta in the individual layers can be seen more clearly in the cross sections (Fig. 1B-B’’’, arrowheads). To test if Inx2 is expressed between the SPG and WG layers, we expressed mCD8::RFP in the SPG and used Nervana 2 endogenously tagged with GFP (Nrv2::GFP) to label the WG. We observed lines of Inx2 puncta (Inx2 plaques) along the SPG (Fig. 1C,D,D’’’, white arrowheads) and WG boundary (Fig. 1D’’). The prominent Inx2 labeling at the SPG-WG boundary suggests that Inx2 plaques are present between the SPG and WG (Fig. 1C, D-D’’’, white arrowheads). Apart from the SPG-WG boundary, Inx2 puncta were also observed in the center of the nerve within the WG membrane labeled with Nrv2::GFP (Fig. 1C, yellow arrowhead). We also analyzed the distribution of Inx1 in peripheral nerves where the SPG and WG were labeled with mCD8::RFP and Nrv2::GFP, respectively. Similar to Inx2 labeling, Inx1 was prominent in the SPG-WG boundary (Fig. 1E, F-F’’, white arrowheads). These findings provide strong evidence that Inx2 is present between all three peripheral glial layers and is expressed along with Inx1 at the SPG-WG boundary as well as within the WG at larval stages. To test if Inx1 and Inx2 were co-localized, we analyzed the distribution of both Inx1 and Inx2 and found that Inx1 and Inx2 puncta co-localize along the SPG-WG boundary (Fig. 1G, H-H’’’, white arrowheads) as well as within the WG membrane (Fig. 1G, yellow arrowhead). Overall, our data suggested that both Inx1 and Inx2 are expressed in all three glial layers and may be constituents of the same junctional complexes.

**Table 1.**
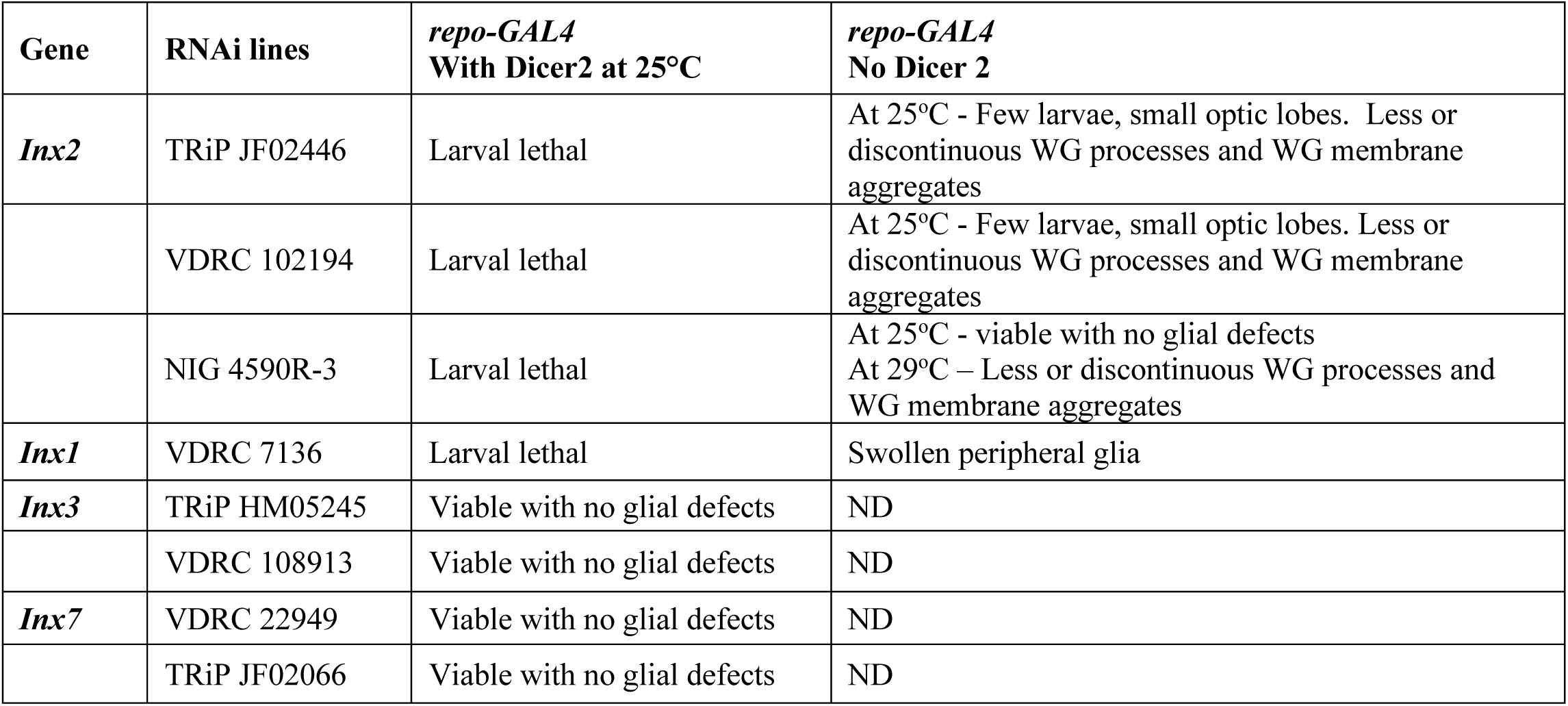
Summary of Inx RNAi phenotypes. Each Inx RNAi was expressed in all glia using the pan glial driver *repo-GAL4* driving mCD8::GFP. Knockdown experiments were conducted in the presence or absence of Dicer2 at the temperatures indicated. Peripheral nerves were screened for glial and nerve defects.

| Gene | RNAi lines | *repo-GAL4*<br>With Dicer2 at 25°C | *repo-GAL4*<br>No Dicer 2 |
| --- | --- | --- | --- |
| *Inx2* | TRiP JF02446 | Larval lethal | At 25°C - Few larvae, small optic lobes. Less or discontinuous WG processes and WG membrane aggregates |
|  | VDRC 102194 | Larval lethal | At 25°C - Few larvae, small optic lobes. Less or discontinuous WG processes and WG membrane aggregates |
|  | NIG 4590R-3 | Larval lethal | At 25°C - viable with no glial defects<br>At 29°C – Less or discontinuous WG processes and WG membrane aggregates |
| *Inx1* | VDRC 7136 | Larval lethal | Swollen peripheral glia |
| *Inx3* | TRiP HM05245 | Viable with no glial defects | ND |
|  | VDRC 108913 | Viable with no glial defects | ND |
| *Inx7* | VDRC 22949 | Viable with no glial defects | ND |
|  | TRiP JF02066 | Viable with no glial defects | ND |

**Figure 1.**
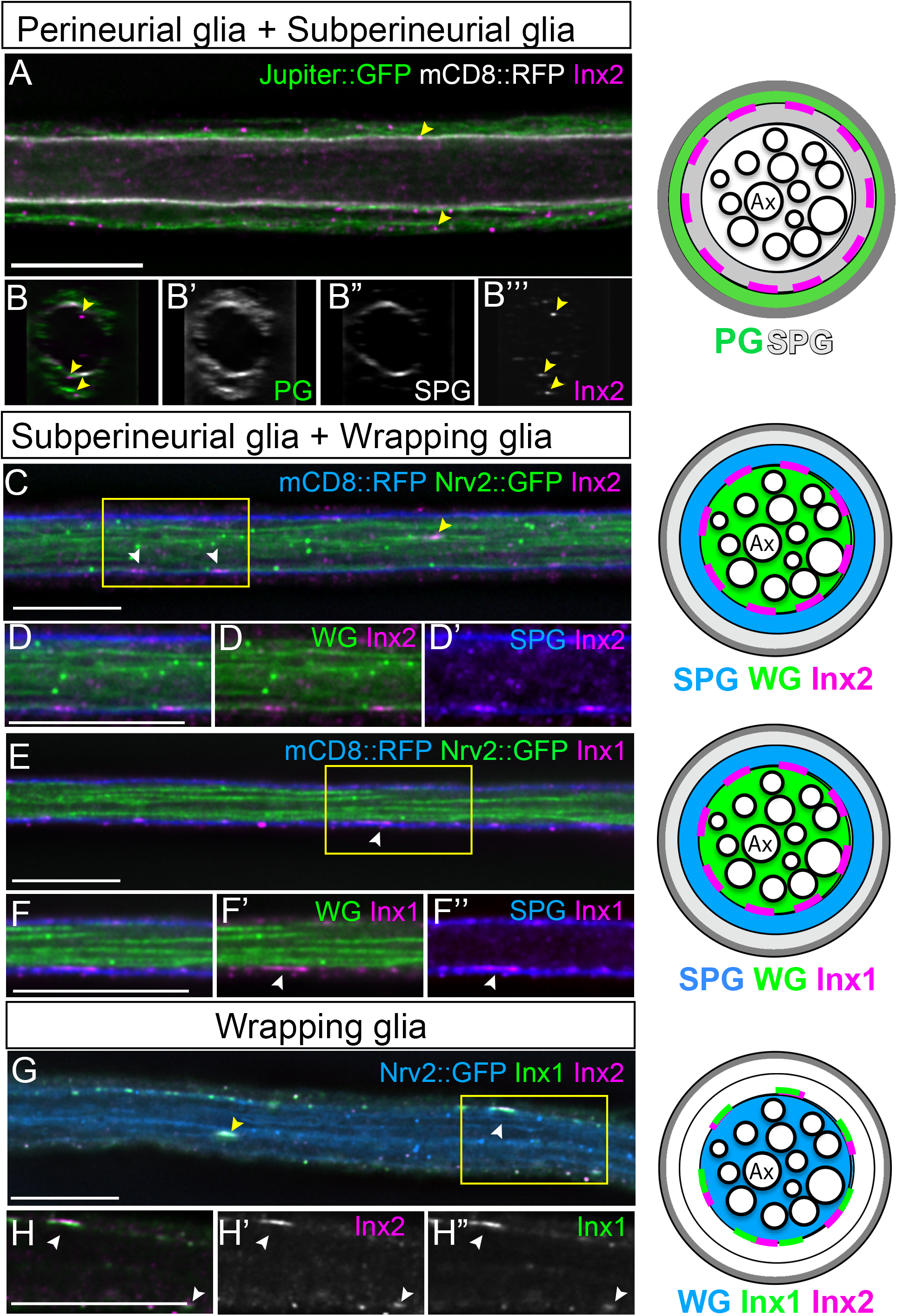
Innexin 1 and Innexin 2 are expressed in the larval peripheral nerve **A, B**: Longitudinal section (A) or cross section (B-B’’’) of a control peripheral nerve with the perineurial glia (PG) membrane and subperineurial glia (SPG) membranes labeled with Jupiter::GFP (green) and mCD8::RFP (white) respectively. Inx2 immunolabeling (magenta) identified puncta (yellow arrowheads) present in the PG and SPG, and within the wrapping glia (WG) in the center of the nerve. **C-D, E-F:** Longitudinal sections of control peripheral nerves with Inx2 (C-D) and Inx1 (E-F) immunolabeling (magenta) in the subperineurial glia and wrapping glia. SPG membrane are labeled with mCD8::RFP (blue), and WG membranes labeled with Nrv2::GFP (green). The yellow boxes were digitally magnified (200x) and shown in D-D’’ and F-F’’ respectively. White arrowheads in C-D and E-F indicate the Inx2 and Inx1 puncta respectively in the SPG. Inx2 puncta in the WG are indicated by the yellow arrowhead (C). Both Inx2 and Inx1 expression puncta are observed in the SPG-WG boundary (C, D’-D’’’ and E,F-F’’’, white arrowheads). **G-H:** Inx1(green) and Inx2 (magenta) form plaques along the SPG-WG boundary (G, H-H’’’, white arrowheads) and within the WG membrane (G, yellow arrowhead). Scale bars: 15μm.

### Loss of Innexin1 and Innexin2 affect the glial cells of the Drosophila larval PNS

We then further investigated the role effects of Inx1 and Inx2 on the peripheral glia using *repo-GAL4* paired with RNAi knockdown, and tested for changes in glial morphology using a fluorescently tagged membrane marker (mCD8::GFP). Inx2 knockdown experiments were carried out using three independent Inx2-RNAi lines with or without Dicer2 to increase the effectiveness of the RNAi (Table 1). All Inx2-RNAi lines with Dicer2 were lethal at early larval stages suggesting that Inx2 is specifically required in glial cells during the early stages of larval development but not during embryogenesis. Without Dicer2 two RNAi lines, Inx2-RNAi (TRiP) and Inx2-RNAi (VDRC), resulted in a larvae surviving to the third instar stage. These larvae had an overall reduction in body size, including smaller brain lobes similar to CNS phenotypes observed previously (Holcroft et al., 2013). We observed reduced and fragmented glial membranes along the length of 67% of nerves (n=3 larvae) with the TRiP RNAi (*repo> Inx2-RNAi TRIP*)(Fig. 2C, D). Control larvae (*repo>*) did not show any glial defects (n=4 larvae)(Fig. 2A,A’ and B,B’). Similar phenotypes were observed using the Inx2 NIG RNAi line (*repo> Inx2-RNAi NIG*), which was less efficient than the others, but with increased expression at 29°C, we observed glial membrane aggregates in 49% of nerves (n=5 larvae) (Fig. 2G,H). Control nerves at 29°C (*repo>*) (n=5 larvae) did not have the fragmented glia phenotype (Fig. 2E,F). For all Inx2 RNAi lines, the glial membrane aggregates were observed in the interior of the nerve and the outer glial membranes appeared normal (Fig. 2 E’,F, G’,H’) suggesting that knockdown of Inx2 in all glia led to defects in WG morphology.

**Figure 2.**
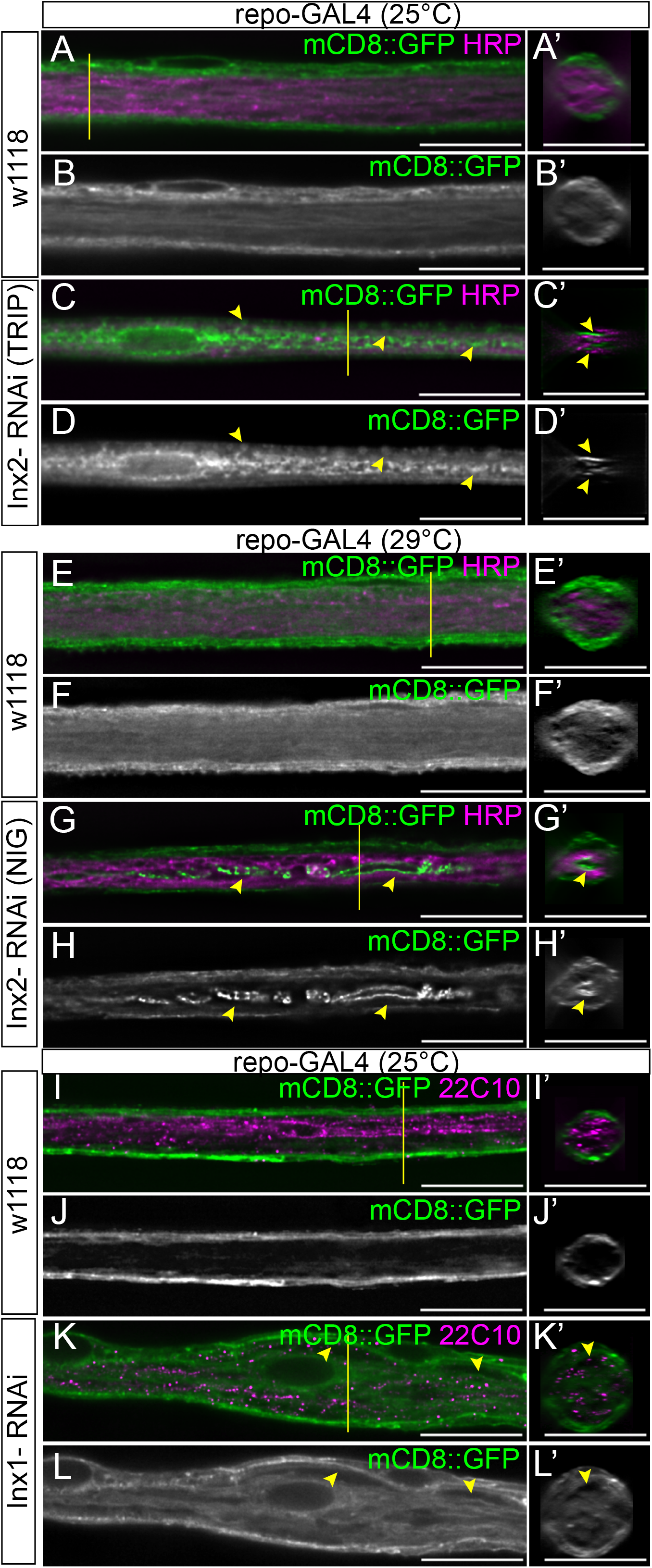
Knockdown of Inx2 in all glia leads to fragmentation of the inner glial membrane, whereas knockdown of Inx1 leads to glial swellings. Longitudinal sections of 3^rd^ instar larval nerves with *repo-GAL4* driving mCD8::GFP (green) to label glial membranes and axons labeled with anti-HRP or anti-22C10 (magenta). The yellow lines indicate the region from which the cross sections were taken. **A-B:** *repo-GAL4* at 25°C. (E-F) Control *repo-GAL4* at 29°C. (I-J) Control *repo-GAL4* at 25°C. In controls the axons (magenta) are present in the center of the nerve and completely surrounded by the glial membrane (green). In the side projections (A’), the PG and SPG membranes surround the entire nerve and the WG membranes fill the core of the nerve **C-D:** *repo>Inx2 RNAi* (TRIP) at 25°C. (G-H) *repo>Inx2 RNAi* (NIG) at 29°C. Peripheral nerves are thinner compared to control nerves and the glial membranes (green) in the center of the nerve were disrupted with membrane fragments (arrowheads, C, G). In the side projections, the outer glial membranes still surround the nerves (C’, G’) but the glial membrane is collapsed or concentrated in the center of the nerve without ensheathing the axons (arrowheads, C’,G’). **K-L:** *repo>Inx1* at 25°C. Swellings were observed between the different glial membranes (green) (yellow arrows, K, L) and the glial membranes surrounded the nerve and filled the center of the nerve (yellow arrows, K’,L’). Scale bars: 15μm.

We next tested the role of Inx1 in peripheral glia. Knockdown of Inx1 in all glia led to smaller brain lobes as observed in previous studies (Holcroft et al., 2013; Speder and Brand, 2014). Furthermore, pan glial knockdown of Inx1 (*repo>Inx1-RNAi*) resulted in peripheral glia swelling in 32% of nerves (n= 12 larvae) (Fig. 2K-H’) compared to control larvae (*repo>mCD8::GFP*), which had no swelling (n=8 larvae)(Fig. 2I-J). Overall, the Inx1 loss of function phenotypes were different from those generated with the Inx2 RNAi, particularly in that Inx2 knockdown did not lead to glial swelling. Our results from the pan-glial knockdown indicate a role for both Inx1 and Inx2 in peripheral glia.

### Innexin1 roles in individual glial layers

Our next step was to investigate the role of each Inx in the individual glial layers using glial layer specific drivers. To test the role of Inx1 in the SPG, we used *Gli-GAL4* to drive RNAi along with mCD8::RFP to mark the SPG membranes and Nrv2::GFP to mark the WG membranes. Inx1 knockdown in SPG (*Gli>Inx1-RNAi, Dicer2*,) did not affect the morphology of the SPG or the neighboring WG, even in the presence of Dicer2 (Fig. 3C-D’; Fig. 4I; Table 2) and the morphology of these glial layers was similar to control larvae (*Gli>Dicer2*)(Fig. 3A-C’; Fig. 4I; Table 2). To test the role of Inx1 in the WG, we knocked down Inx1 using the *Nrv2-GAL4* driver and mCD8::GFP to mark the membranes. Co-expressing Dicer2 with the Inx1-RNAi at 25°C (*Nrv2>Inx1-RNAi, Dicer2)* resulted in 86% of nerves with reduced and discontinuous WG strands, where neighboring WG processes failed to meet (Fig. 3G-H’; Fig. 4I; Table 2). Control larvae expressing Dicer2 at 25°C alone (*Nrv2>Dicer2)* had no WG defects Fig. 3E-F’; Fig. 4I; Table 2). To test the role of Inx1 in the outermost perineurial glia, we used the 46F-GAL4 driver and a membrane marker, mCD8::RFP. Inx1 knockdown (*46F>Inx1-RNAi*) did not affect the morphology of the PG (n=5 larvae) (Fig. 3K-L’) which were comparable to control (*46F>)*(n=4 larvae)(Fig. 3I-J’). The Inx1 knockdown results suggest that Inx1 has a function in the WG but not in the SPG or PG (or maybe redundant with Inx2 in these outer glial layers). Moreover, knockdown of Inx1 in each of the three glial layers alone did not result in glial swellings observed with knockdown in all glia.

**Table 2.** Summary of screen Inx2 knockdown phenotypes in the WG Each driver paired with UAS-mCD8::RFP was crossed to the transgenes listed. Controls were drivers plus mCD8::RFP crossed with *w[1118]*. In part 2, the number of affected nerves included both nerves with reduced or single WG processes and nerves with membrane fragments. Experiments were conducted at 25°C.

| Driver | UAS-transgenes | larvae | % nerves with WG phenotypes |  |  |
| --- | --- | --- | --- | --- | --- |
| *Part 1* |  |  | wild type | reduced or single processes | membrane fragments |
| **SPG**<br>*Gli-GAL4*<br>*UAS-mCD8::RFP* | *control* | n=11 | 100% | 0% | 0% |
|  | *Dicer2* | n=7 | 100% | 0% | 0% |
|  | *Inx2-RNAi* | n=18 | 34% | 57% | 11% |
|  | *Inx2-RNAi, Dicer2* | n=11 | 19% | 47% | 34% |
|  | *Inx1-RNAi* | n=6 | 100% | 0% | 0% |
|  | *Inx1-RNAi, Dicer2* | n=5 | 100% | 0% | 0% |
| **WG**<br>*Nrv2-GAL4*<br>*UAS-mCD8::RFP* | *control* | n=6 | 98% | 2% | 0% |
|  | *Dicer2* | n=6 | 100% | 0% | 0% |
|  | *Inx2-RNAi* | n=23 | 3% | 79% | 24% |
|  | *Inx2-RNAi, Dicer2* | n=9 | 0% | 77% | 23% |
|  | *Inx1-RNAi* | n=5 | 15% | 85% | 0% |
|  | *Inx1-RNAi, Dicer2* | n=5 | 14% | 86% | 0% |
| *Part 2* |  |  | wild type | affected |  |
| **WG**<br>*Nrv2-GAL4*<br>*UAS-mCD8::RFP* | *p35* | n=5 | 87.2% | 11.8% |  |
|  | *Inx2-RNAi, p35* | n=16 | 7.5% | 91.5% |  |
|  | *Inx2-RNAi, lacZ* | n=9 | 15.8% | 84.2% |  |
|  | *Atg1-RNAi* GD7149, *Inx2-RNAi* | n=5 | 14.6% | 85.4% |  |
|  | *Atg1-RNAi* VSH330433, *Inx2-RNAi* | n=6 | 11.4% | 88.6% |  |
|  | *Atg18-RNAi, Inx2-RNAi* | n=7 | 19.1% | 81.0 % |  |

**Figure 3.**
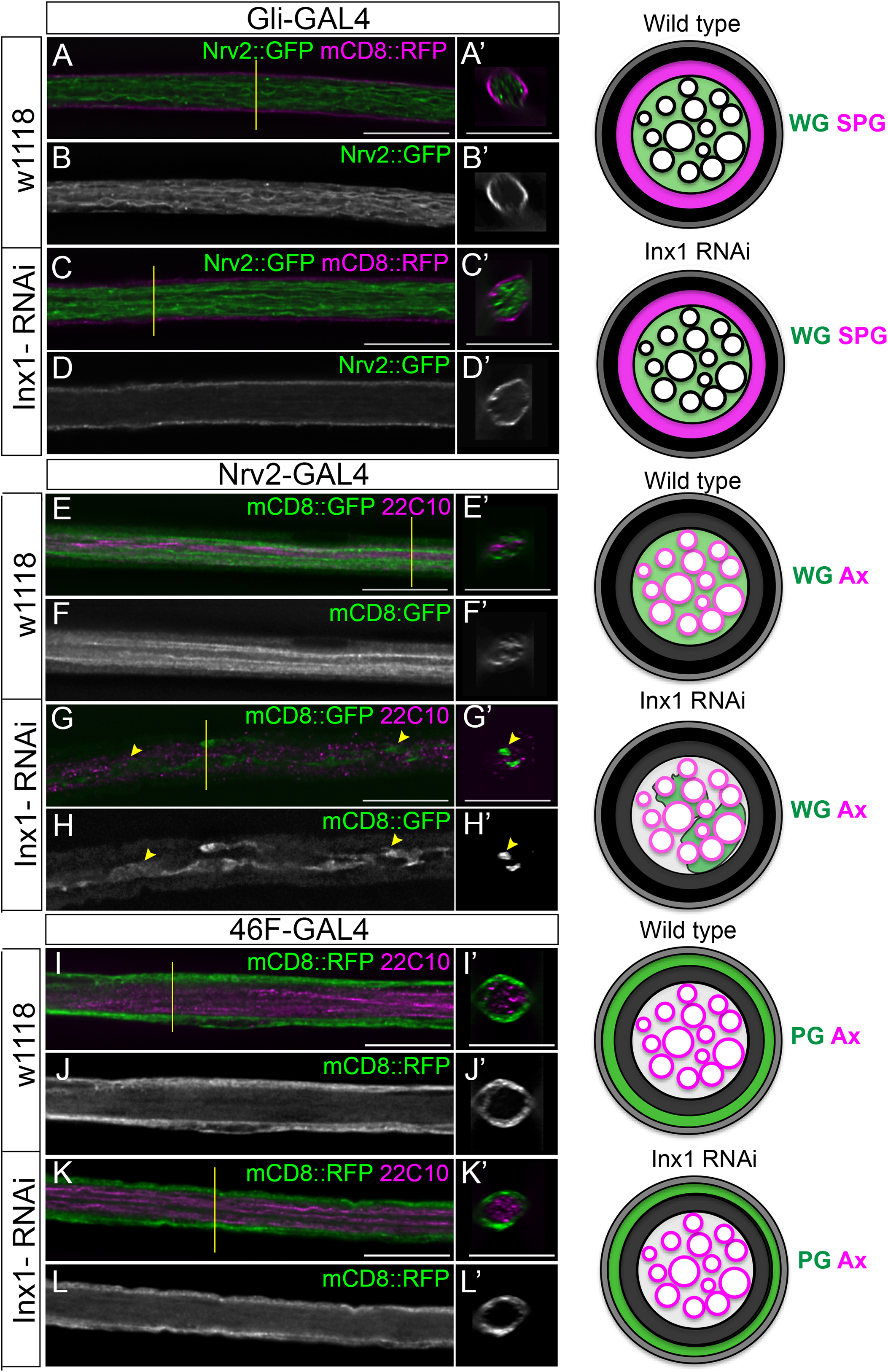
Knockdown of Inx1 in the wrapping glia leads to defects but knockdown in the other glial layers does not affect morphology. Nerves for 3^rd^ instar larvae in longitudinal and cross sections. The yellow lines indicate the region from which the cross sections were taken. A schematic representation of the labeled glial layers in control and Inx1 knockdown nerves are shown to the right with each glial layer and axons (Ax) indicated. **A-D:** Subperineurial glia. (A-B) Control - *Gli-GAL4*. (C-D) *Gli>Inx1-RNAi*. SPG membranes labeled with mCD8::RFP (magenta) and WG membranes labelled with Nrv2::GFP (green). The SPG and WG membranes in the *Gli>Inx1-RNAi* nerve were similar to the control nerve. WG extends processes along (A,C) and throughout the core of the nerve (A’,C’). The thin SPG membrane flanks the nerve (A,C) surrounding the WG (A’,C’). **E-H:** Wrapping glia. (A-B) Control - *Nrv2-GAL4*. (C-D) *Nrv2>Inx1-RNAi*. WG membranes were labeled with mCD8::RFP (green) and axons immunolabeled with 22C10 (magenta). Strands of WG wrap (green) around axons (magenta) in the control nerve (E) and cross sections (E’). WG strands were reduced and discontinuous in *Nrv2>Inx1-RNAi* nerves (G, yellow arrowhead) and the WG membrane does not fully wrap around axons in some regions of the nerve (G’, yellow arrowhead). **I-L:** Perineurial glia. (I-J) Control – *46F-GAL4*. (K-L) *46F>Inx1-RNAi*. Nerves with PG membranes labeled with mCD8::RFP (green) and axons labeled with 22C10 (magenta). PG membranes surround the entire nerve in both control and *46F>Inx1-RNAi*. Scale bars: 15μm.

**Figure 4.**
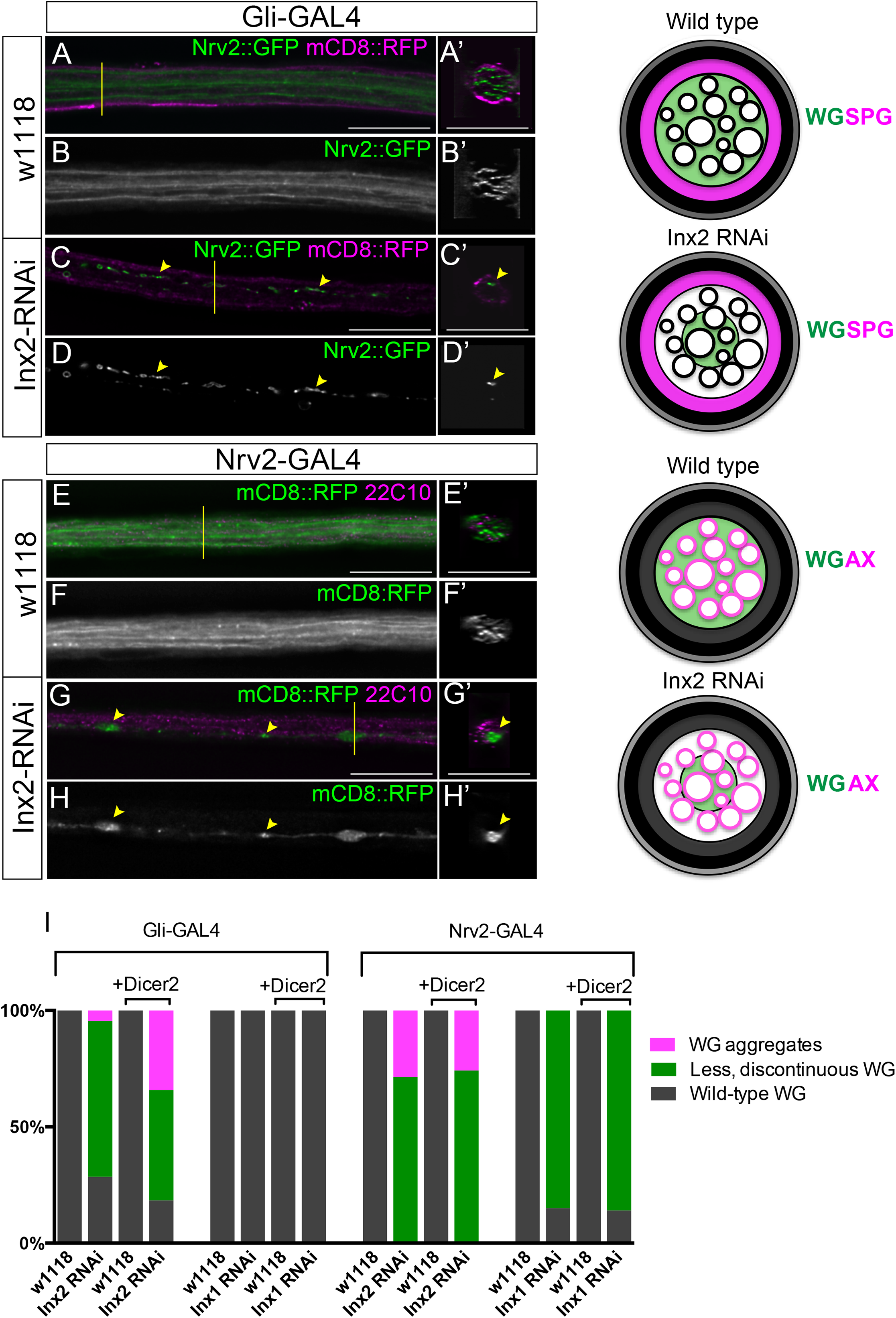
Knockdown of Inx2 in subperineurial glia or wrapping glia leads to fragmentation of the wrapping glia. Nerves for 3^rd^ instar larvae in longitudinal and cross sections. The yellow lines indicate the region from which the cross sections were taken. A schematic representation of the labeled glial layers in control and Inx1 knockdown nerves are shown to the right with each glial layer and axons (Ax) indicated. **A-D:** Subperineurial glia (SPG). (A-B) Control – *Gli-GAL4* (A-B) and *Gli>Inx2-RNAi* (C-D). Nerves with SPG and wrapping glia membranes labeled with mCD8::RFP (magenta) and Nrv2::GFP (green), respectively. The WG (green) in control nerves extend processes along the entire length of the nerve and the thin SPG membrane (magenta) surrounds the WG (A’). WG membranes wrap around the peripheral axons (A, A’). WG membrane aggregates (green, C) are present along the length of the nerve in *Gli>Inx2-RNAi*. Remnants of the WG membrane (green) are found in the center of the nerve (C’, arrowhead) and do not wrap around axons (magenta) in *Gli>Inx2-RNAi*. **E-H:** Wrapping glia (WG). Control (E) and *Nrv2>Inx2-RNAi* (G) peripheral nerves with WG membranes labeled with mCD8::RFP (green) and axons immunolabeled with 22C10 (magenta). The yellow lines indicate the region from which the cross sections were taken. Several strands of WG wrap (green) around axons (magenta) in the control peripheral nerve (E, E’) and cross sections indicate that WG membrane surrounds axons (magenta). Loss of WG strands in *Nrv2>Inx2-RNAi*, with only a single WG strand (green) present in the peripheral nerve (G,G’). WG membrane in the *Nrv2>Inx2-RNAi* nerve is only present at the center with an uneven morphology and does not wrap around axons (magenta). **I:** Comparison of the WG phenotypes observed when Inx2 and Inx1 were knocked down in the SPG and the WG. The WG phenotypes were divided into three categories: wild type WG (grey); less and discontinuous WG strands (green); WG aggregates (magenta). The specific percentage of nerves that fall under the three categories of WG phenotypes are detailed in Table 2.

### Innexin2 roles in individual glial layers

To determine if Inx2 is required in one or all three glial layers, we knocked down Inx2 in individual glial layers using the most efficient Inx2-RNAi (TRiP) line. For knockdown in the perineurial glia, we used the 46F-GAL4 driver and labeled the PG membrane with mCD8::RFP and the WG with Nrv2::GFP. We did not observe any phenotypes in any glial layer (*46F>Inx2-RNAi)* (n=6 larvae, data not shown). To test the role of Inx2 in the SPG, we used *Gli-GAL4* with mCD8::RFP to mark the SPG membranes and Nrv2::GFP to mark the WG. Knockdown of Inx2 (*Gli>Inx2-RNAi)* had no effect on the morphology of the SPG membrane (Fig. 4C). Surprisingly, the knockdown of Inx2 in the SPG did affect the morphology of the WG, where we observed a range of WG phenotypes (Fig. 4C-D’,I; Table 2). 57% of the nerves had either fewer WG strands or a single, discontinuous WG membrane. WG fragments, similar to those seen in the pan glial knockdown of Inx2, were observed in 11% of nerves. Control larvae (*Gli>*) did not show any WG defects (Fig. 4A-B’,I; Table 2). Thus, the loss of Inx2 from the SPG had little or no effect on the SPG, but rather had a cell non-autonomous effect on the WG.

To analyze the role of Inx2 in the WG, we knocked down Inx2 using the *Nrv2-GAL4* driver and mCD8::RFP to mark the membranes (*Nrv2>Inx2-RNAi*). We observed single strands of WG processes in 79% of nerves (Fig. 4G-H’,I; Table 2) compared to the multiple strands normally observed in the WG (Fig. 5A,B). Membrane aggregates in the WG were observed in 24% of nerves, while control larvae (*Nrv2>*) (Fig. 4E-F’,I; Table 2) did not show any glial defects. Our results suggest a role for Inx2 within the WG and the SPG to ensure proper WG ensheathment of the peripheral axons.

**Figure 5.**
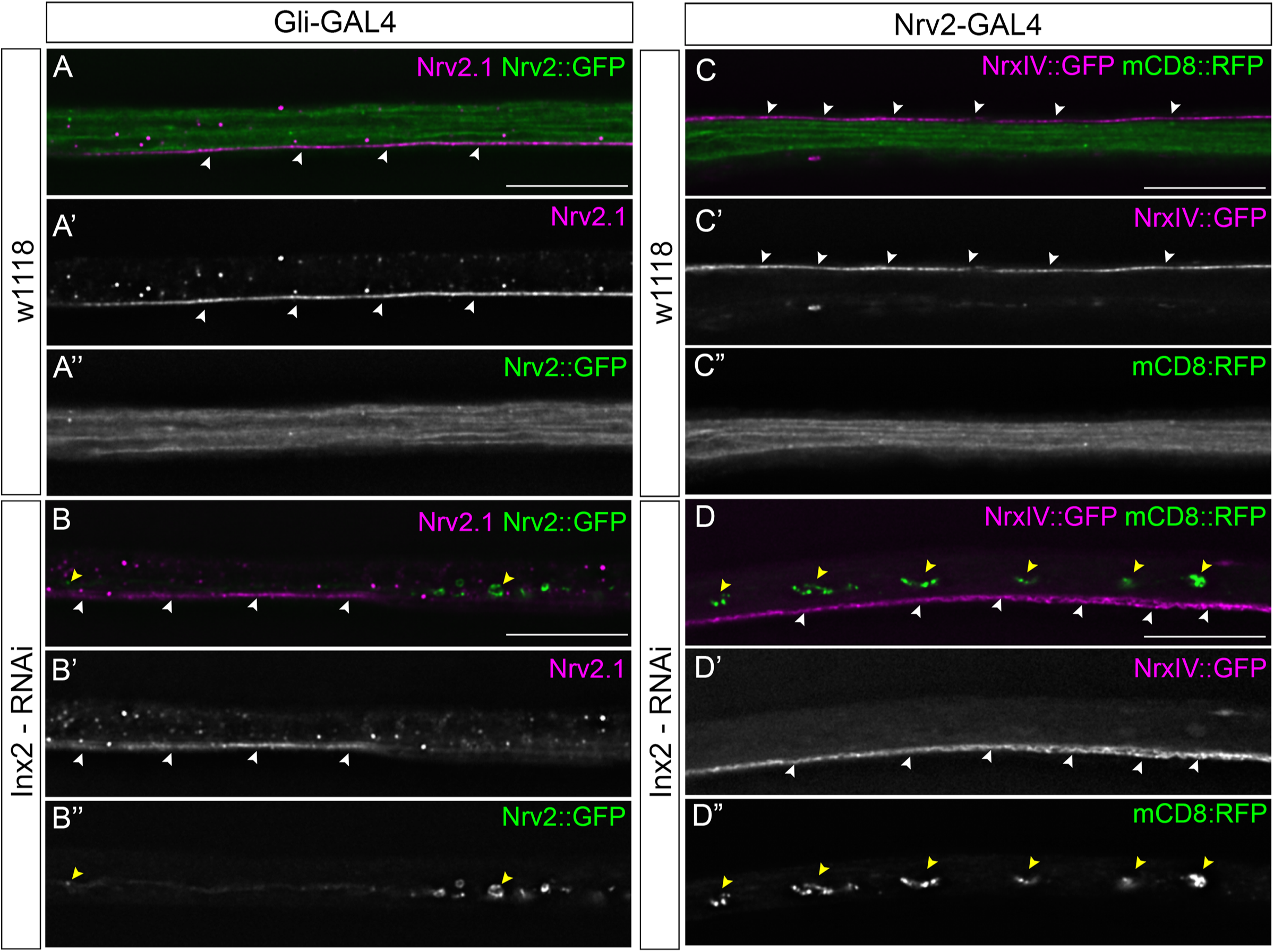
Knockdown of Inx2 in the SPG or WG does not affect septate junction morphology. **A-B:** Control (*Gli-GAL4*) (A) and *Gli>Inx2-RNAi* (B) peripheral nerves with WG membranes labeled with Nrv2::GFP (green) and the SJ domain labeled with a Nrv2.1 antibody (magenta). **C-D:** Control (*Nrv2-GAL4*) (C) and *Nrv2>Inx2-*RNAi (D) peripheral nerves with WG membranes labeled with mCD8::RFP (green) and the SJ domain labeled with NrxIV::GFP (magenta). WG (green) in the control nerves (A”, C”) extend processes along the entire length of the nerve and the SJ domain (magenta) is continuous along the length of the nerve (A’,C’, white arrowheads). With Inx2 knockdown WG membrane aggregates (B”, D”, yellow arrowheads) are present along the length of the nerve in Gli>Inx2-RNAi (B) and Nrv2>Inx2-RNAi (D). SJ morphology is not affected (B’, D’, white arrowheads)

To further investigate if the loss of Inx2 affected the morphology of the SPG layer, we analyzed the distribution of two key septate junction proteins (NrxIV::GFP and Nrv2.1) when Inx2 was knockdown in either the SPG (Fig. 5A-B) or the WG (Fig. 5C-D). Normal, continuous WG strands were observed in control nerves (Fig. 5A”, C”) and the SJ markers formed a single junctional domain along the length of the nerve (Fig. 5A’,C’) (n=4 larvae). Knockdown of Inx2 in either the SPG or WG resulted in loss of WG strands and fragmentation (Fig. 5B”,D”) but did not affect the distribution or levels of NrxIV::GFP or Nrv2.1. Overall Inx2 appears to be key for the morphology and wrapping of the WG but not for the formation of the SPG or septate junction domain.

### Innexin2 knockdown in WG affects processes and ensheathment of axons

We next wanted to characterize in greater detail the phenotypes we observed within the WG when Inx2 was knocked down in the SPG. Using both fluorescence and transmission electron microscopy, a range of phenotypes were observed in the WG as a result of Inx2 knockdown in the SPG (*Gli>Inx2-RNAi*) compared to control larvae (*Gli-GAL4*) (Fig. 6). At the ultrastructural level, we found that the overall structure of the nerve was intact in both control (Fig. 6A’’) and *Gli>Inx2-RNAi* larvae (Fig. 6B’’-E’’). Overall, SPG morphology appeared normal at the ultrastructural level with SJs present along the SPG layer as expected (Fig. 6A’’-E’’, magenta arrowheads). In the control nerves, WG strands in the light microscope images (Fig. 6A,A’) correspond to the processes of a single WG cell, which at the TEM level ensheath individual or bundles of axons (Fig. 6A’’). While the SPG appeared intact and normal, knockdown of Inx2 in the SPG resulted in a range of WG phenotypes. At the ultrastructural level, we observed less WG processes and the extent of axonal ensheathment was reduced compared to the control (Fig. 6B’’-E’’). In the light microscope images, some nerves had clear breaks within the WG membrane and reduced WG strands around axons (Fig. 6B, B’). In other nerves, the WG failed to ensheath the majority of the axons and appeared as individual processes though the core of the nerve (Fig. 6C, C’). In some nerves, few WG processes extended along the length of the nerve but were interrupted by breaks in the membrane (Fig. 6D,D’). In other nerves, only a few fragments were observed at the distal end of a single WG process (Fig 6E,E’). Therefore, knock down of Inx2 in the SPG resulted in a failure to ensheath axons and often fragmented WG processes.

**Figure 6.**
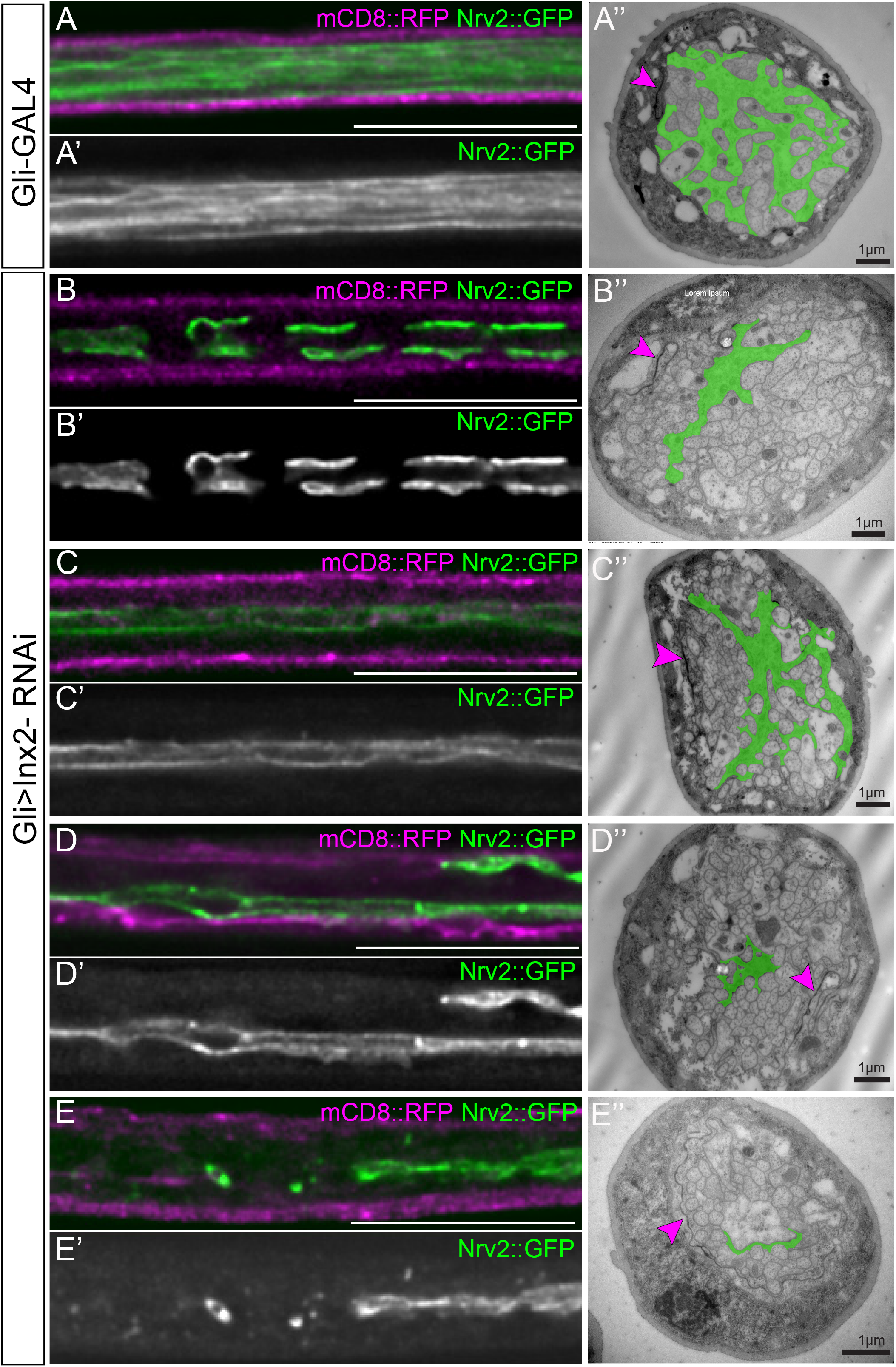
Knockdown of Inx2 in the subperineurial glia results in a range of wrapping glia phenotypes. 3^rd^ instar peripheral nerves with SPG and WG membranes labeled with mCD8::RFP (magenta) and Nrv2::GFP (green) respectively with ultrastructural images with the WG false-coloured green (A”-E”) to the right. **A:** Control (*Gli-GAL4*). WG strands (green) are present and extend along the length of the control peripheral nerve. (A”) In cross section, WG (green) extend processes and wrap around axons or bundles of axons. **B-E:** *Gli>Inx2-RNAi*. Loss of Inx2 in the SPG results in a range of WG phenotypes, with nerves showing wrapping glia membrane aggregates (B), reduced wrapping glia strands (C), discontinuous strands (D) or strands ending in fragments (E). Representative TEM sections that correspond to the WG phenotypes with reduced strands (B”,C”) or reduction to single strands (D”, E”). Septate junctions are indicated in the TEM sections (A’’-E’’, magenta arrowheads). Scale bars: 15μm (A-E), 1 μm (A”-E”)

Of note, the WG fragments were only observed within the nerve extension region (NER), which is wrapped by both the first and second WG, and never in the muscle field area (MFA) which is wrapped by the third WG (Fig. 8A). When we mapped the location of the WG phenotypes within each peripheral nerve, we found that majority of the WG phenotypes with knockdown in both the SPG (n=14 larvae) and WG (n=10 larvae) were observed within the NER at the point where the processes of the first WG contact those of the second WG. Taken together these results suggest that the loss of Inx2 in SPG or the WG result in a range of WG defects indicating that some level of contact or communication between the two glial layers (SPG and WG) is required for normal WG morphology and axonal ensheathment and that loss of Inx2 leads to destabilization of the WG wrap and WG contact.

### Innexin2 is required in the SPG for WG integrity but not WG survival

The loss of the WG wrap and the presence of membrane fragments could be due to a number of possible mechanisms including cell death or loss of trophic or nutritive support. To determine if the WG aggregates in *Nrv2>Inx2-RNAi* larvae are due to apoptosis, we expressed p35, a baculoviral protein that blocks caspase activation (Hay et al., 1994). Even in the presence of p35, WG aggregates and loss of WG processes were observed with Inx2-RNAi (Table 2). In addition, the WG in *Gli>Inx2-RNAi* and *Nrv2>Inx2-RNAi* larvae were not positive for Drosophila caspase-1 (Dcp-1), an effector caspase in apoptosis (Song et al., 1997) (n=14; data not shown) further supporting that disruption of the WG is not due to apoptosis.

Another means of membrane fragmentation could be through autophagy. We reduced autophagy by co-expressing RNAi lines to knockdown Atg1 and Atg18 with Inx2-RNAi (*Nrv2>Atg1-RNAi, Inx2-RNAi & Nrv2>Atg18-RNAi, Inx2-RNAi*) and compared to control where LacZ was co-expressed with Inx2-RNAi (*Nrv2>lacZ, Inx2-RNAi*). Knockdown of autophagy proteins did not reduce the percentage of Inx2-RNAi induced WG defects (Table 2) compared to control indicating that autophagy does not play a role. To further test for autophagy we expressed the tandem GFP-mCherry-Atg8a marker in the WG while simultaneously knocking down Inx2 (*Nrv2> GFP-mCherry-Atg8a, Inx2-RNAi*). The tandem *GFP-mCherry-Atg8a* marker co-labels the non-acidified components with both GFP and mCherry signals while labeling the acidic late endosomes or lysosomes with mCherry as GFP is lost in the acidic environment. We observed no differences between control (*Nrv2> GFP-mCherry-Atg8a*, n=8 larvae) and Inx2 knockdown nerves (*Nrv2> GFP-mCherry-Atg8a,Inx2-RNAi*) where the majority of the signals were yellow suggesting Inx2 in the WG does not lead to increase in autophagic flux (data not shown). The absence of apoptotic and autophagy makers along with failure to rescue the WG defects associated with Inx2 knockdown by blocking apoptosis and autophagy point towards a different mechanism in mediating changes to the WG morphology.

### Knockdown of Innexins in the SPG and the WG leads to locomotion defects

We next tested whether the loss of Inx2 affected nervous system function and specifically the motility of 3^rd^ instar larvae. We found that larvae with knockdown of Inx2 in the SPG (*Gli>Inx2-RNAi*) (n=211) travelled less and moved at a significantly slower speed than control larvae (*Gli-Gal4*) (n=86)(Fig. 7A,B). Similarly, larvae with Inx2 knockdown in the WG (*Nrv2>Inx2-RNAi*) (n=121) were significantly slower and travelled less distance than control (*Nrv2-Gal4*) larvae (n=157) (Fig. 7A.B). Larvae of both genotypes (*Gli>Inx2-RNAi* and *Nrv2>Inx2-RNAi*) failed to eclose and were lethal at pupal stages, suggesting that the continued expression of Inx2 in the SPG and WG is necessary for survival.

**Figure 7.**
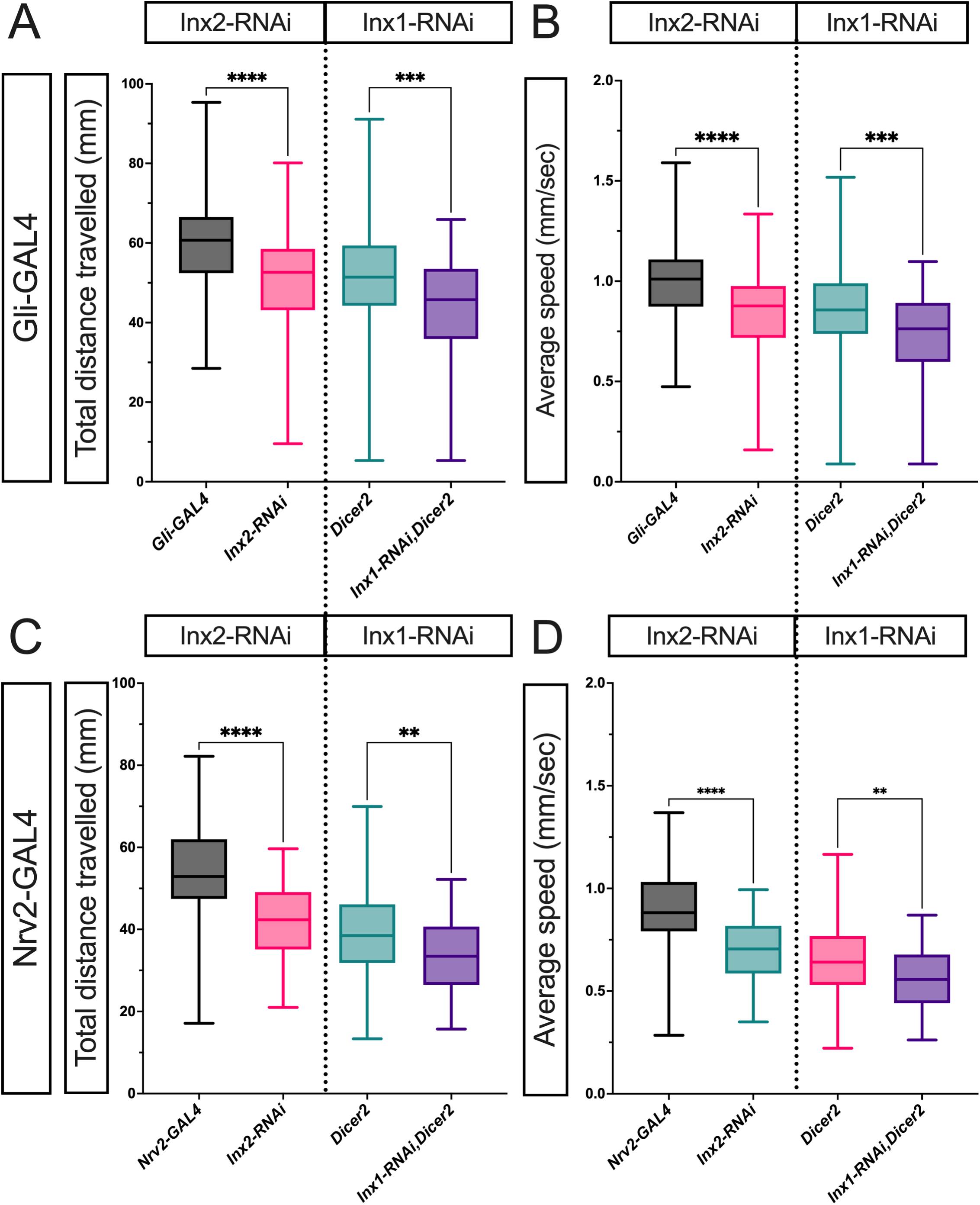
Inx knockdown in the subperineurial and wrapping glia affects larval locomotion. Total distance travelled and average speed were determined for 3^rd^ instar larvae. Boxes indicate the 25^th^ to 75^th^ percentiles with the median indicated. The whiskers indicate the minimum to maximum values. **A-B**: Subperineurial glia (*Gli-GAL4*). The total distance travelled (A) or average speed (B) in SPG control (*Gli>)*(n=86) compared to *Gli>Inx2-RNAi* (n=211) and *Gli>Dicer2*(n=226) compared to *Gli>Inx1,Dicer2* (n=69). **C-D:** Wrapping glia (*Nrv2-GAL4*). The total distance travelled (C or average speed (D) in WG control (*Nrv2>*)(n=157) compared to *Nrv2>Inx2-RNAi* (n=121) and *Nrv2>Dicer2*)(n=133) compared to *Nrv2>Inx2-RNAi* (n=50). Statistical significance was determined by a one-way ANOVA with Tukey’s multiple comparisons test for each driver (*Gli>, Gli>Inx2-RNAi, Gli>Dicer2, Gli>Inx1-RNAi, Dicer2*) and (*Nrv2>, Nrv2>Inx2-RNAi, Nrv2>Dicer2, Nrv2>Inx1-RNAi,Dicer2*). A: ****p<0.0001, ***p=0.0001; B: ****p>0.0001, ***p=0.0001. C: ****p<0.0001, **p=0.0037; D: ****p<0.0001, **p<0.0036

To test Inx1, we knocked down Inx1 in the SPG and tested larval locomotion. We found that *Gli>Inx1-RNA, Dicer2* larvae (n=69) had significantly slower speeds than control larvae (*Gli>Dicer2*) (n=226)((Fig. 7C,D). Knockdown of Inx1 in the WG (*Nrv2> Inx1-RNAi, Dicer2*) (n=50) also significantly affected locomotion and distance travelled compared to control (*Nrv2>Dicer2*) larvae (n=133)(Fig. 7C,D). Inx1 and Inx2 loss in the SPG and WG both affect larval locomotion however this might reflect a secondary effect of loss of Innexin in CNS given that both drivers express within CNS glial populations.

### Inx2 based gap junctions mediate calcium pulses in the peripheral SPG but not WG

Having established that Inx1 and 2 are present and have a function in peripheral glia, we next wanted to test for channel function of Inx-based junctions in the PNS. In the CNS, synchronous calcium oscillations in the SPG are dependent on heteromeric Inx1/Inx2 gap junctions (Speder and Brand, 2014) and Ca^2+^ signals are propagated between perineurial glia using Inx2 containing gap junctions (Weiss et al., 2022). But whether gap junctions are also involved in propagation of calcium pulses in the peripheral SPG or WG had not been previously tested. We first tested if there were Ca^2+^ signals within the peripheral glia using the GFP Calcium indicator GCaMP6S (Chen et al., 2013). GCaMP6S was expressed in the SPG (*Gli>GCaMP6S*) and imaged in live intact 3^rd^ instar larvae. In controls, we observed changes in Ca^2+^ levels in the first peripheral SPG (Fig. 8A,A’-B,B’), which represents the first of four SPG cells that covers each peripheral nerve. SPG1 covers the Nerve Extension Region (NER), whereas the remaining SPG nuclei are present in the muscle field area (MFA)(von Hilchen et al., 2013). We found that the changes in Ca^2+^ initiated in the ventral nerve cord presumably in the SPG of the CNS and occured in the first peripheral SPG as a pulse (Fig. 8B,B’) rather than a wave (n= 9 larvae) (Fig 3A,A’, Movie 1). The calcium pulses were not observed in all nerves in and in total occurred in 49.3% of nerves, particularly in the nerves of abdominal segments A6-A8 at the posterior end of the ventral nerve cord. There was a range of responses where some nerves had no detectable changes in Ca^2+^, others pulsed infrequently, and others pulsed multiple times.

**Figure 8.**
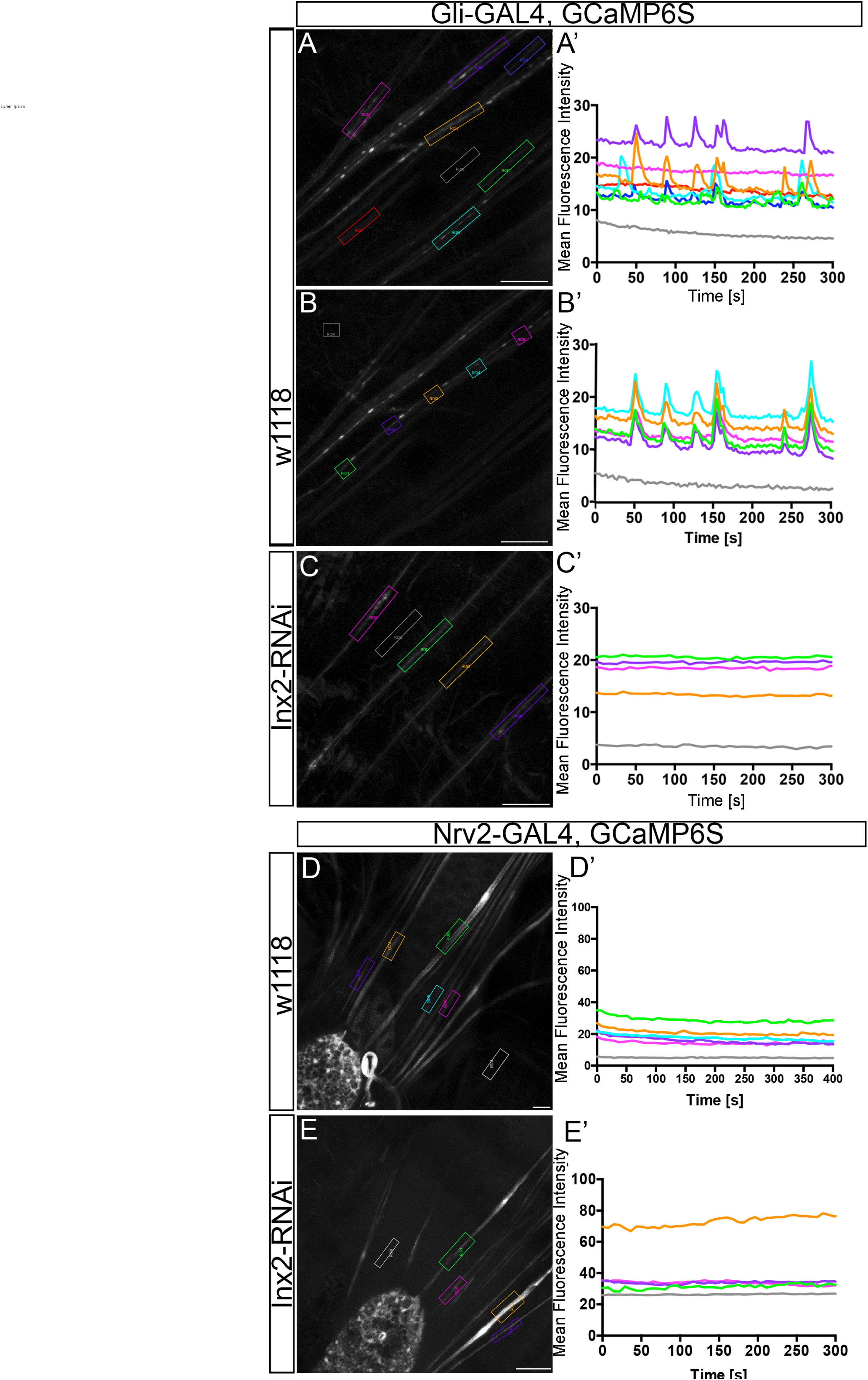
Calcium pulses are confined to the subperineurial glia and require Inx2. 3^rd^ instar larval nerves in live preparations with GCaMP6S expressed in subperineurial (A-C) or wrapping glia (D-E). The change in mean fluorescence intensity over time (seconds) is plotted in the graphs (A’-E’) indicated by the regions of interest (ROIs). ROI 18 in each image (grey, A-E) was placed away from the peripheral nerve and represents the basal GCaMP6S signal. **A-C:** Subperineurial glia. GCaMP6S driven by Gli-GAL4 in control (*Gli>GCaMP6S*) (A-B’) or with Inx2-RNAi (*Gli>GCaMP6S, Inx2-RNAi*) (C’C’). (A,A’) Calcium pulses were present in some but not all control nerves (ROIs: 1, 12-14,16; green, purple, orange, light blue and blue, respectively). (B,B’) Control nerve where the ROIs are placed along the same nerve. The change in mean fluorescence intensity occur as a pulse along the peripheral nerve. (C) Knockdown of Inx2 in the SPG (*Gli>Inx2-RNAi, GCaMP6S*) blocked the pulses with no observed changes in mean fluorescence intensity. **D-E**: Wrapping glia. GCaMP6S driven by Nrv2-GAL4 in control (*Nrv2>GCamp6S*) (D) or with Inx2-RNAi (*Nrv2>GCaMP6C, Inx2-RNAi*) (E). Calcium pulses were not observed in the peripheral nerves (ROIs:1-5; green, purple, orange, light blue and pink, respectively) of control (D) or Inx2-RNAi expressing WG (E). The ROI 18 (grey, D,E) were placed in a region where peripheral nerves were absent to measure basal level of the GCaMP6S signal. Scale bars: 25 μm

When Inx2 was knockdown in the SPG, calcium pulses were not observed in any peripheral nerves (*Gli>Inx2-RNAi, GCaMP6S*) (n=10 larvae) (Fig. 8C,C’, Movie 2). This suggests that Inx2-based gap junction channels mediate calcium signals in the SPG. Since Inx1 and Inx2 form heteromeric channels in the CNS SPG (Speder and Brand, 2014), we tested the role of Inx1 in mediating the peripheral Ca^2+^ pulses. Knockdown of Inx1 in the SPG (*Gli>GCaMP6S, Inx1-RNAi*) did not disrupt the calcium pulses in the peripheral nerves (n=5 larvae) even in the presence of Dicer 2 (*Gli>GCaMP6S, Inx1-RNAi, Dicer2*) (n=5 larvae) and we observed calcium pulses in a pattern and frequency similar to control larvae (*Gli>GCaMP6S*) (data not shown). This suggests that calcium signals in peripheral SPG cells are gap junction mediated and that Inx2 may function as an independent homomeric channel as observed previously (Holcroft et al., 2013).

We then used the same imaging conditions to record calcium in the peripheral WG by expressing GCaMP6 alone (*Nrv2>GCaMP6S*) as well as with Inx2-RNAi (*Nrv2>GCaMP6S, Inx2-RNAi*). Calcium pulses were not observed in any WG (n= 7 larvae) (Fig. 8D, Movie 3) and the knockdown of Inx2 had no effect (*Nrv2>GCaMP6S, Inx2-RNAi*) (n= 7 larvae) (Fig. 8E, Movie 4). We did observe microdomain calcium oscillations in the glia of the ventral nerve cord suggesting the GCaMP6S was expressed at sufficient levels for Ca^2+^ detection. Thus, while there were robust Inx2-dependent Ca^2+^ pulses observed in the SPG, Ca+2 pulses were absent from the neighbouring WG suggesting that functional gap junctions do not form between the SPG and WG.

### Inx2 has channel-independent functions between the SPG and WG

Inx2 could mediate SPG to WG communication as a cell adhesion protein, given that gap junctions can also function as cell adhesion proteins in a channel independent manner (Bruzzone et al., 1996; Kumar and Gilula, 1996; Phelan et al., 1998; Elias et al., 2007). To gather a better understanding of Inx2 function, we utilized a dominant negative transgene with RFP fused to the N-terminus of Inx2 (RFP::Inx2), which leads to a loss of Inx2 function (Nakagawa et al., 2010; Speder and Brand, 2014; Oshima et al., 2016; Sahu et al., 2017). Expression of the dominant negative RFP::Inx2 in the SPG (*Nrv2::GFP, Gli>RFP::Inx2*) resulted in a range of WG phenotypes similar to Inx2 RNAi (Fig. 9D-I’). WG fragments were observed in 11% of the nerves (Fig. 9D-F’) while 100% of the nerves had discontinuous strands or reduced numbers of strands (Fig. 9G-I’). We observed no differences in the SPG. Control larvae using a C-terminally tagged Inx2 expressed in the SPG did not have any WG phenotypes (*Nrv2::GFP, Gli>Inx2::RFP*)(n=5 larvae) (Fig. 9A-C’). Next, we expressed the dominant negative RFP::Inx2 in the WG (*Nrv2>RFP::Inx2*) and found that expressing RFP::Inx2 in the WG generated discontinuous WG strands and a failure to wrap (Fig. 9M-R’). Moreover, the WG discontinuities were observed in the NER where the first and second WG contact each other (data not shown), similar to our prior observation in Inx2 knockdown. Overall, the dominant negative Inx2 generated the same phenotypes that we observed with our Inx2-RNAi knockdown.

**Figure 9.**
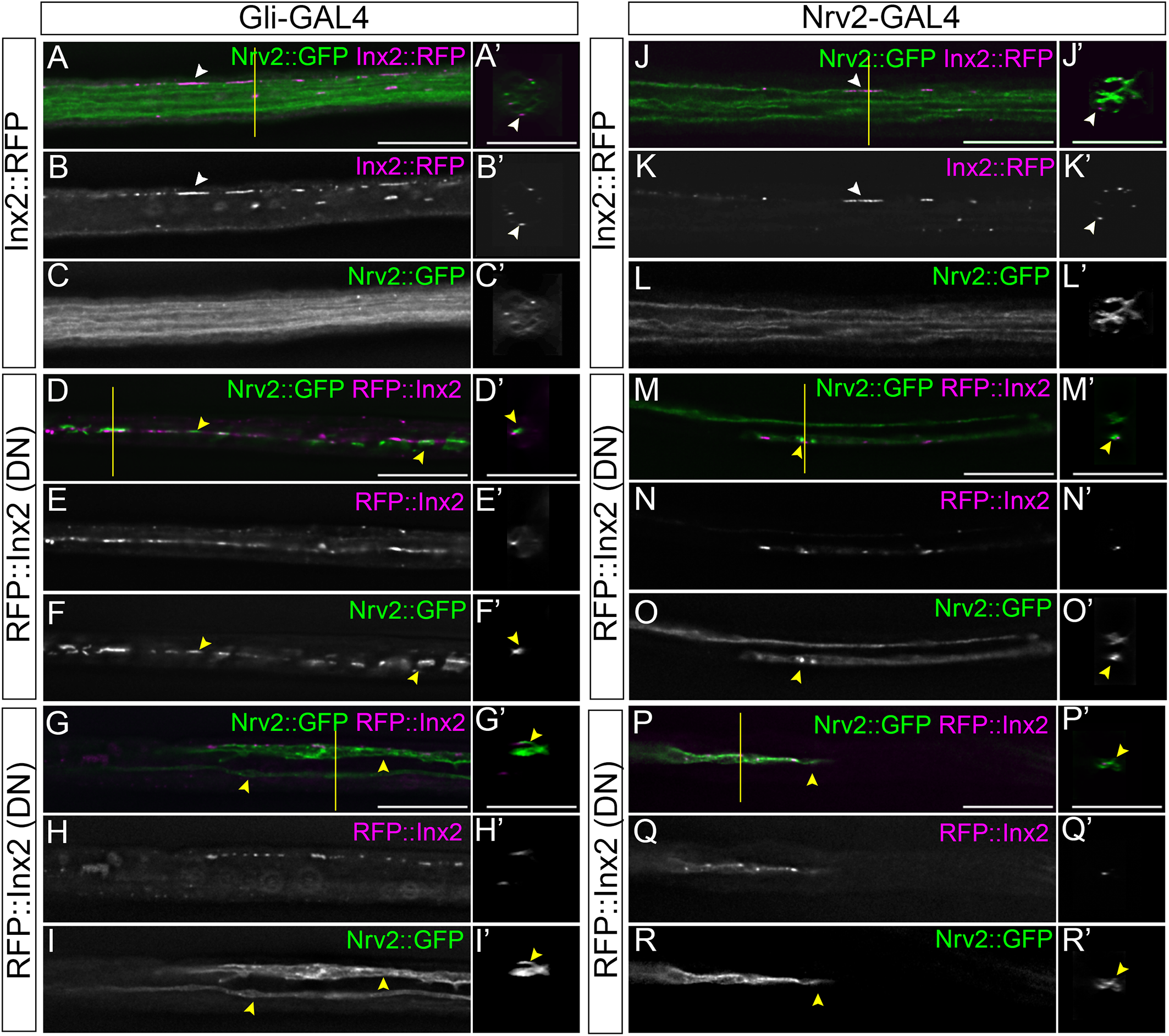
Dominant negative Inx2 triggers the same WG phenotypes as Inx2 RNAi **A-C:** *Gli>Inx2::RFP* control nerves expressing control Inx2::RFP in the SPG (magenta) and WG membranes labeled with Nrv2::GFP (green). The WG membrane in the control nerve (A, green) extends its processes along the entire length of the nerve and Inx2::RFP plaques (magenta, white arrowheads) are present in the neighboring SPG (A, magenta). **D-I:** *Gli>RFP::Inx2* nerves with the dominant negative Inx2 mutant (RFP::Inx2, magenta) driven in the SPG and the WG membranes labeled with Nrv2::GFP (green). WG membrane aggregates (green, yellow arrowheads) are present along the length of some nerves in Gli>RFP::Inx2 larvae (D,F) whereas others have discontinuous WG membranes (G,I). RFP::Inx2 (magenta) is localized to the SPG-WG boundary (D-E’, G-H’). **J-L**: *Nrv2>Inx2::RFP* control nerves expressing Inx2::RFP (magenta) and mCD8::GFP (green) in the WG. The WG membrane (green) extends processes along the entire length of the nerve and Inx2::RFP plaques (magenta, white arrowheads) are present in the WG, along the SPG-WG boundary. **M-R:** *Nrv2>RFP::Inx2* peripheral nerves with the dominant negative Inx2 mutant (RFP::Inx2, magenta) driven in the WG along with mCD8::GFP (green). Single, discontinuous WG strands (green, yellow arrowheads) were observed (M-O) with others nerves containing disrupted WG strands (P-R). Agregates of WG membrane were found in the center of the nerve (M,M’). The yellow lines indicate the region from which the cross sections were taken. Scale bars: 15μm.

To determine if the RFP::Inx2 in the SPG integrated into junctions linking the SPG and WG membranes, we examined the distribution of RFP::Inx2 in relation to Inx1 across different focal planes to capture the interface between the WG and SPG. In control nerves expressing Inx2 with C-terminal RFP in the SPG (*Nrv2::GFP, Gli>Inx2::RFP*), 100% of the Inx2::RFP plaques were co-localized with Inx1 plaques in the SPG (Fig. 10A,A’, yellow arrowheads, z=39) and formed Inx1/2 plaques that consistently corresponded with Inx1 positive plaques in the opposing WG membrane (Fig. 10B,B’, green arrowheads)(n=5 nerves)(Fig. 10E). When the dominant negative RFP::Inx2 was expressed in the SPG (*Nrv2::GFP, Gli>RFP::Inx2*), 96.7% of the Inx2 plaques colocalized with Inx1 in the SPG (Fig. 10C,C’, yellow arrowheads)(n=9 nerves). However only 3.7% of the dominant negative RFP::Inx2 /Inx1 plaques in the SPG corresponded with Inx1 in the opposing WG (Fig. 10D,D’,white arrowheads)(Fig. 10E)(Movie 5). This observation suggests that the dominant negative RFP::Inx2 is able to integrate into hemichannels with Inx1 but that these hemichannels in the SPG are unable to connect with those in the WG.

**Figure 10.**
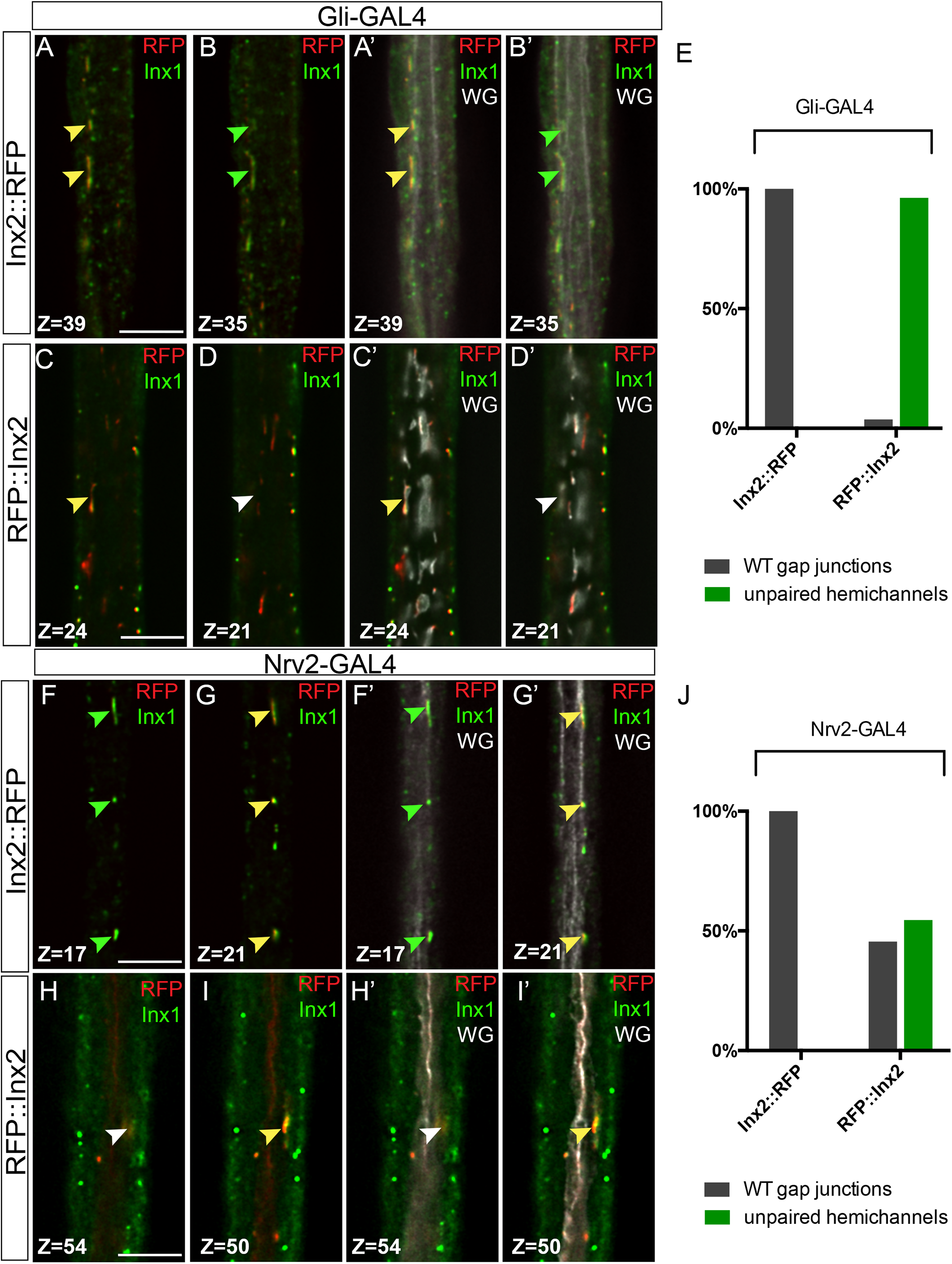
Dominant negative Inx2 blocks coupling with Inx1 in the neighbouring glia. 3^rd^ instar nerves with Inx2::RFP or RFP::Inx2 (red) expressed in the SPG or WG with Inx1 immunolabeled (green). Individual Z-stacks are indicated a two focal planes to capture the association of Inx2 with Inx1 (first two panels) and in the context of the WG membrane (Nrv2::GFP, gray) (second two panels. Scale bars: 7.5μm. **A-B:** *Gli>Inx2::RFP* control nerves where Inx2::RFP (red) is expressed in the SPG with Inx1 immunolabelled (green). Inx2::RFP integrates with Inx1 in the SPG (yellow plaques)(**A**,**A’**, yellow arrowheads) that correspond to Inx1 immunolabeling (green plaques) in the underlying WG (B,B’, green arrowheads). WG processes (Nrv2::GFP, grey) extend along the entire length of the nerve (A’,B’). **C-D**: *Gli>RFP::Inx2* nerves with dominant negative Inx2 mutant (*RFP::Inx2*) driven in the SPG with Inx1 immunolabelling (green). RFP::Inx2 integrates with Inx1 in the SPG (yellow plaques) (C,C’, yellow arrowhead) but the absence of Inx1 in the underlying WG (D,D’, white arrowhead) suggests a lack of coupling between the SPG and WG (C’,D’). **E:** The percentage of wild type gap junctions (grey) and unpaired hemichannels (green) observed in control (*Gli>Inx2::RFP*, n=3 nerves) compared to nerves expressing dominant negative Inx2 in the SPG (*Gli>RFP::Inx2*, n=9 nerves). **F-G:** *Nrv2>Inx2::RFP* control nerves with Inx2::RFP (red) expressed in the WG (Nrv2::GFP, gray) and Inx1 immunolabelled (green). Inx2::RFP integrates with Inx1 in the WG (yellow plaque)(G,G’, yellow arrowheads) and corresponds to Inx1 the overlying SPG (green plaque)(F,F’, green arrowheads. WG processes extend along the entire length of the nerve (G’). **H-I**: *Nrv2>RFP::Inx2* nerves with the dominant negative Inx2 mutant (*RFP::Inx2*, red) driven in the WG and Inx1 immunolabelled (green). WG membranes were labelled with Nrv2::GFP (grey; **H’**,**I’**). The dominant negative RFP::Inx2 integrates with Inx1 in the WG (yellow plaques)(I,I’, yellow arrowhead) but the absence of Inx1 (green) in the corresponding area in the SPG (H,H’, white arrowhead) suggests a lack of coupling between the hemichannels in the SPG and WG (H’,I’). Scale bars: 7.5μm **J:** The percentage of wild type gap junctions (grey) and unpaired hemichannels (green) observed in control (*Nrv2>Inx2::RFP*, n= 2 nerves) compared to nerves expressing dominant negative Inx2 in the WG (*Nrv2>RFP::Inx2*, n=7 nerves).

We carried out a similar analysis with expression in the WG. In control nerves expressing C-terminal tagged Inx2::RFP (*Nrv2 >Inx2::RFP*), plaques containing Inx1 and Inx2 (Fig. 10F,F’, yellow arrowheads) were observed along the SPG-WG boundary. 100% of the observed Inx1/Inx2 plaques in the WG were coupled with Inx1 positive plaques in the SPG (Fig. 10F-F’, green arrowheads; Fig. 10J) (n=3 nerves). In WG expressing dominant negative RFP::Inx2 (*Nrv2 >RFP::Inx2*) (n=7 nerves), Inx1/Inx2 plaques were observed in the WG strands or fragments (Fig. 10H,H’, yellow arrowheads) but only 45.5% coupled with Inx1 positive plaques in the overlying SPG (Fig. 10H,H’, white arrowheads; Fig. 10J). These results suggest that the dominant negative RFP::Inx2 interferes with the ability of the gap junctions to form contacts between the WG and the SPG.

Our next step was to further test the function of Inx2 either as a channel or as an adhesion protein. Miao et al. (2020) generated Inx2 transgenes that allow these functions to be separately tested. The Inx2[L35W] mutant eliminates channel activity by replacing a highly conserved lysine 35 to tryptophan but leaves channel-independent functions intact (Depriest et al., 2011; Baker et al., 2013). The Inx2[C265S] mutant blocks all Innexin functions through the loss of a highly conserved extracellular Cys residue. Both mutants plus a wild-type Inx2 were tagged at the C-terminus with RFP and are resistant to the Inx2 TRIP RNAi. We crossed each to the SPG driver (Moody-GAL4) with WG labeled with Nrv2::GFP to assess the ability of each to rescue the Inx2-RNAi mediated WG phenotypes. Inx2 knockdown in the control (*Nrv2::GFP, Moody>Inx2-RNAi, mCD8::RFP*) (n=16) generated the expected WG phenotypes (Fig. 11A,E). Wildtype Inx2::RFP (*Nrv2::GFP, Moody>Inx2-RNAi, R-Inx2::RFP*) (n=11) rescued the WG defects (Fig. 11B,E) as did the Inx2[L35W]::RFP mutant (*Nrv2::GFP, Moody>Inx2-RNAi, R-Inx2 [L35W]::RFP*)(n=18)(Fig. 11C,E). The Inx2[C256S]::RFP mutant (*Nrv2::GFP, Moody>Inx2-RNAi, R-Inx2[C256S]::RFP*) (n=15) was not able to rescue (Fig. 11D,E). Overall, our results point to an adhesion function for Inx2 between the SPG and WG membranes.

**Figure 11.**
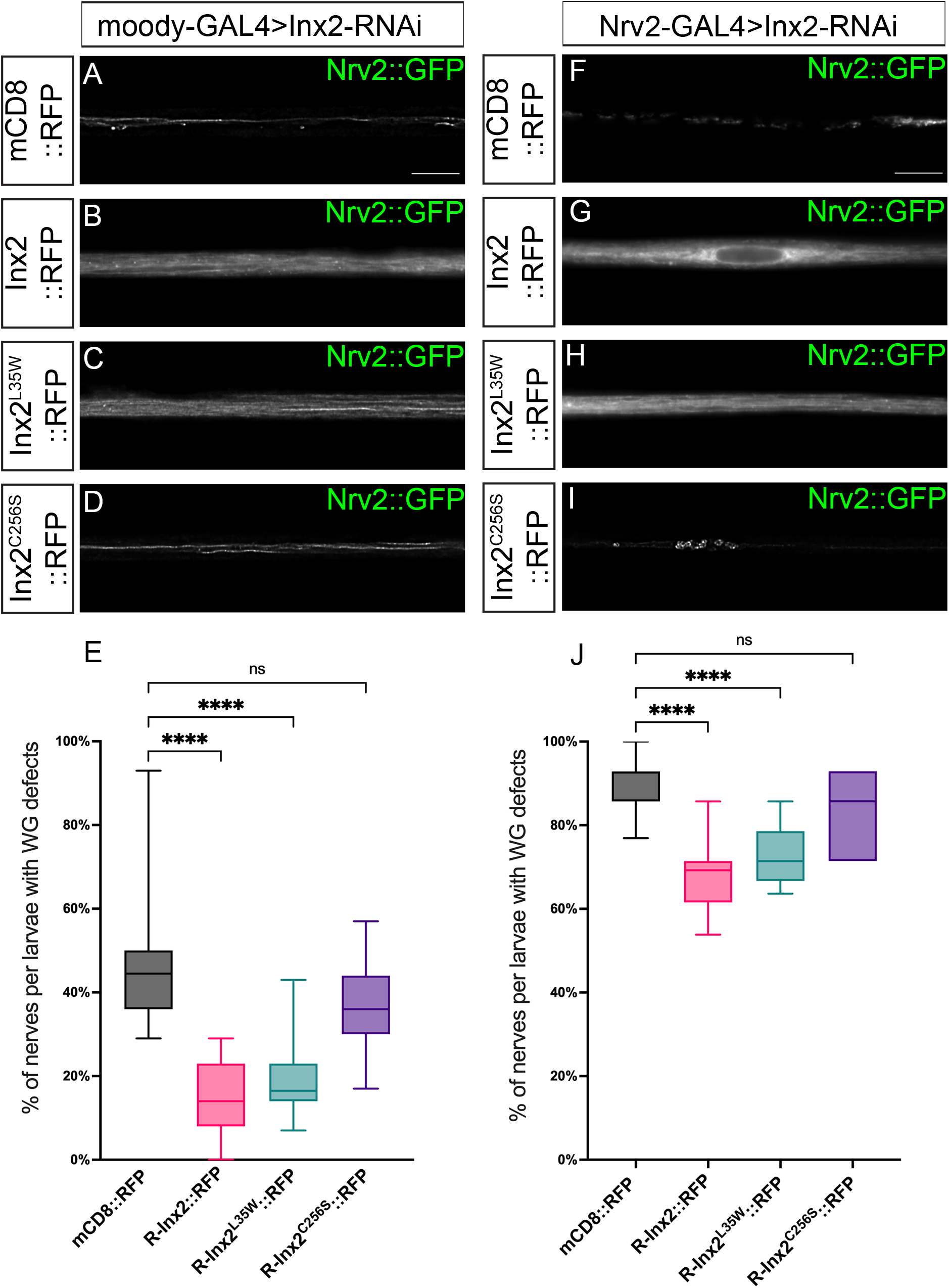
Rescue of the Inx2-RNAi phenotype by both wild type and a channel-deficient Inx2 mutant 3^rd^ instar nerves with the WG membrane marked by Nrv2::GFP. Scale bars: 15μm. **A-E**: Subperineurial glia. *Moody-GAL4* driven Inx2-RNAi (*moody>Inx2-RNAi*) with (A) mCD8::RFP,(B) Inx2::RFP, (C) Inx2::RFP[L35W], (D) Inx2::RFP[C256S]. Inx2-RNAi lead to the disruption of the WG and these phenotypes were rescued by Inx2::RFP and Inx2::RFP[L35W] but not Inx2::RFP[C256WS]. (E) The degree of rescue was quantified and statistical significance was determined by a one-way ANOVA with Tukey’s multiple comparisons test. Boxes indicate the 25^th^ to 75^th^ percentiles with the median indicated. The whiskers indicate the minimum to maximum values. ****p<0.0001, ns=not significant. **F-J**: Wrapping glia. *Nrv2-GAL4* driven Inx2-RNAi (*Nrv2>Inx2-RNAi*) with (F) mCD8::RFP, (G) Inx2::RFP, (H) Inx2::RFP[L35W], (I) Inx2::RFP[C256S]. Inx2-RNAi lead to the disruption of the WG and these phenotypes were rescued by Inx2::RFP and Inx2::RFP[L35W] but not Inx2::RFP[C256WS]. (J) The degree of rescue was quantified and statistical significance was determined by a one-way ANOVA with Tukey’s multiple comparisons test. Boxes indicate the 25^th^ to 75^th^ percentiles with the median indicated. The whiskers indicate the minimum to maximum values. ****p<0.0001, ns=not significant.

We then tested the ability of the Inx2 mutants to rescue when expressed in the wrapping glia. Using Nrv2-GAL4, we observed that both wild-type Inx2 (*Nrv2::GFP, Nrv2>Inx2-RNAi, R-Inx2::RFP*)(n=11) and Inx2[L35W] (*Nrv2::GFP, Nrv2>Inx2-RNAi, R-Inx2 [L35W]::RFP*)(n=11) partially rescued the wrapping glial ensheathment phenotypes (Fig. 11G,H,J). On the other hand, Inx2[C256S] (*Nrv2::GFP, Nrv2>Inx2-RNAi, R-Inx2[C256S]::RFP*)(n=11) had the same percentage of WG defects as control (*Nrv2::GFP, Nrv2>Inx2-RNAi, CD8::RFP*)(n=12) (Fig. 11F,I,J). Both Inx2 wildtype and Inx2[L35W] also reduced the severity of the Inx2-RNAi phenotypes, where with control and Inx2[C256S] 25% and 34% of the nerves had the WG fragmentation phenotype respectively. This is compared to 8% with wild-type Inx2 and 0% with Inx2[L35W]. Overall, our rescue experiments point to a channel-independent function of Inx2. Inx2 likely functions as an adhesion protein where loss of cell-cell adhesion disrupts the ability of WG to maintain contact with the SPG or disrupts the WG-WG junctions leading to failure of process extension or stabilization.

## DISCUSSION

The presence and requirement for gap junctions in myelinating Schwann cells (SCs) of the vertebrate PNS is well established. Apposing membranes of the myelin sheath are coupled by autotypic gap junctions to mediate rapid intercellular communication over the myelin layers (Balice-Gordon et al., 1998; Nualart-Marti et al., 2013). In comparison, very little is known about gap junctions in non-myelinating SCs. We show that in *Drosophila*, at least two gap junction proteins, Inx1 and Inx2, are expressed in all the peripheral glial layers. Inx1 and Inx2 were colocalized through the glial layers with clear heteromeric plaques identified between the SPG and WG membranes. Knockdown of Inx2 specifically in WG resulted in failure of the WG to ensheath the peripheral axons and resulted in glial membrane fragments. WG were similarly disrupted when Inx2 (but not Inx1) was knocked down in the SPG suggesting that Inx2 mediates SPG to WG communication and likely acts as an adhesion protein. Consistent with prior work in the vertebrate CNS, which identified gap junctions between oligodendrocytes and astrocytes (Nagy and Rash, 2003; Orthmann-Murphy et al., 2008), we found that Inx2 and Inx2 gap junctions form between two glial layers in the peripheral nervous system, the subperineurial glia (SPG) and wrapping glia (WG).

### Inx1/Inx2 form heteromeric gap junctions in the SPG and WG

Inx1 is known to function in a complex with Inx2 and RNAi knockdown of Inx1 or Inx2 in all glia leads to changes in Ca^2+^ signaling in the CNS and reduction brain lobe size (Holcroft et al., 2013; Speder and Brand, 2014). However studies using Xenopus oocytes found that Inx1 does not form homomeric hemichannels or channels and must pair with other innexins to form functional gap junctions (Holcroft et al., 2013). Conversely Inx2 is able to form homotypic gap junctions albeit with altered properties (Holcroft et al., 2013). We hypothesize that Inx1 and Inx2 form heteromeric channels in both the SPG and the WG, and in the absence of Inx1, Inx2 is present in the form homotypic channels. There are number of observations in support of this model. Only the loss of Inx2 blocked the Ca+2 pulses we observed in the SPG, suggesting that in the Inx1 knockdown experiments homomeric Inx2 channels were able to form and function. Loss of Inx2 but not Inx1 in the SPG resulted in defects in the neighbouring WG suggesting that in the absence of Inx1, Inx2 is still able to meditate SPG to WG adhesion. Loss of Inx1 in the WG lead to far milder phenotypes and never generated the membrane fragments we observed with loss of Inx2. Finally, we observed Inx1/Inx2 heteromeric plaques in close apposition between the SPG and the WG and these were disrupted when a dominant negative Inx2 was expressed in either the SPG or WG. It is possible that Inx1 or Inx2 could form junctions with other Innexins, however it is unlikely to be Inx3 or Inx7 as knockdown did not affect peripheral glial or nerve morphology and knockdown of Inx3 in all glia does not lead to a reduced VNC or brain lobes in larvae (Holcroft et al., 2013). Thus, our data points to Inx1/Inx2 heteromeric channels between the SPG and WG and possibly Inx2 homomeric junctions between SPG-SPG.

### Innexins have two different functions with peripheral glia

We determined that there are two different functions for Innexins within peripheral nerves. In the SPG alone, Innexins appear to mediate a gap junction function as loss of Inx2 resulted in the absence of the Ca+2 pulses we observed in the SPG. This is similar to changes observed in the perineurial glia (PG) where loss of Inx2 blocked Ca2+ signaling between neighbouring PG cells (Weiss et al., 2022). Loss of both Inx1 and Inx2 in the SPG disrupted larval locomotion suggesting a physiological function of the gap junctions in the SPG but this could also be due to disruption of gap junctions in the CNS as both GAL4 drivers are expressed in the CNS. Loss of Inx2 or Inx1 had no effect on PG or SPG morphology and within the SPG did not disrupt the formation of the septate junctions that form the glial blood-nerve barrier suggesting a physiological role for either Inx1 and Inx2 within the outer glial layers of the peripheral nerve and the CNS.

The other function of Innexins in the SPG and WG appears to be channel independent. Many gap junctions have channel-independent functions including Connexins in vertebrate and Innexins (Dbouk et al.,2009; Elias and Kriegstein, 2008; Kameritsch et al., 2013; Leo-Macias et al., 2016; Miao et al., 2020; Wei et al., 2004; Zhou and Jiang, 2014). We hypothesize that the heteromeric junctions formed between the SPG and WG do not function as gap junction channels but rather as an adhesion complex. We observed extensive Inx1/Inx2 plaques between the SPG and WG, yet were unable to detect any Ca+2 pulses within the WG. We were able to disrupt WG morphology and axon wrapping through the loss of Inx2 or the expression of a dominant negative Inx2 within the SPG. Finally, we were able to rescue the Inx2 knockdown WG phenotypes using a transgene that has no channel function but still retains the ability to adhere.

In the context of an adhesion function, knocking down Inx2 in either the SPG or the WG lead to a range of WG phenotypes that included reduced or single WG strands and WG fragments. Loss of Inx1 in the WG but not the SPG generated reduced or single WG strands but no WG fragments. Our results mirror the observations of Kottmeier at al. (2020), who found changes to WG morphology following both Inx1 and Inx2 knockdown and hypothesized that loss of Inx leads to poorly differentiated WG. Similar to our observations, knockdown of Inx1 did result in a mild locomotion defect in 3^rd^ instar larvae and likely due to a decrease in the total distance traveled due to increased curling (Kottmeier et al., 2020). We also observe locomotion defects when Inx2 was reduced in the WG, though as the WG driver also has CNS expression this could be due to a disruption of CNS glial function.

Normally the WG generate an elaborate network of strands that surround single or bundles of peripheral axons and two WG cover the entire extent of the nerve extension region of each nerve. Stabilization of this network likely requires adhesion between the WG and axons, between the WG processes themselves and finally between the WG and overlying SPG. Loss of Inx2 in either glial type leads to a reduction in these strands and the degree of axon wrapping suggesting that Inx2 functions on both sides of the membrane to stabilize the WG ensheathment. With respect to the WG fragments, the frequency was stronger when Inx2 was knocked down in the WG and the majority of fragments were observed in the region where WG1 contacts WG2 in the nerve extension region (Matzat et al., 2015). It is possible that this region is more susceptible to loss of Inx2 and that Inx2 is also necessary to ensure WG to WG adhesion.

Another possible means of WG disruption could be due to changes in the glial process extension and stabilization due to changes in the cytoskeleton and in particular microtubules. Vertebrate connexins (Cx26 and Cx43) can drive migration of neural progenitors (Elias et al., 2007). Cx43 binds to tubulin directly (Giepmans et al., 2001) and both Cx43 and Cx26 stabilize microtubule-rich leading processes in neurons migrating on radial glia (Elias et al., 2007). Drosophila Innexins drive border cell migration where Inx2 and Inx3 are required in border cells and Inx4 within the germline (Miao et al., 2020). All three Innexins regulate microtubules to brace the cells against the morphogenetic forces exerted on the oocyte and border cells. These observations have led to a model where gap-junction mediated microtubule stabilization might contribute to stabilization of cell-cell contacts and thus the loss of WG integrity could be due to changes in the underlying cytoskeleton.

One other means by which Innexins could stabilize the WG is by creating autotypic junctions in the WG as the membrane processes wrap individual or bundles of axons. Connexins form autotypic gap junctions in Schwann cells between the layers of non-compact myelin (Meier et al., 2004) as well as at paranodal loops and Schmidt-Lanterman incisures (Scherer et al., 1995; Spray and Dermietzel, 1995). In myelinating glia, gap junctions are thought to provide a direct radial route for transport of water, ions and small molecules between cytoplasmic myelin layers and given the elaborate processes of the WG the same could be true in Drosophila WG. However, we found no evidence of changes in cell survival or induction of apoptosis or autophagy with the loss of Inx2 in the WG and tests of other metabolic pathways did not affect WG morphology (data not shown). Rather our data supports a model by which Inx1/Inx2 heteromeric channels link the SPG and WG membranes and Inx2 in this complex mediates the adhesion properties of this complex. Rather than forming gap junction channels this complex functions to maintain adhesion between the two glial layers to ensure that the WG processes are stabilized and axon ensheathment and glial-glial adhesion is maintained.

## ACKNOWLEDGMENTS

We thank Drs. Guy Tanentzapf and Reinhard Bauer for the Inx2 and Inx1 antibodies, respectively and Drs. Denise Montell and Andrea Brand for fly stocks. We also thank Dr. Michael Gordon for use their facility to conduct the calcium imaging experiments. This research was funded by a grant from the Natural Sciences and Engineering Research Council of Canada (NSERC) and the Canadian Institutes of Health Research (CIHR).

